# Topoisomerase 2β-dependent nuclear DNA damage shapes extracellular growth factor responses through AKT phosphorylation dynamics to control virus latency

**DOI:** 10.1101/518373

**Authors:** Hui-Lan Hu, Lora A. Shiflett, Mariko Kobayashi, Moses V. Chao, Angus C. Wilson, Ian Mohr, Tony T. Huang

**Affiliations:** Department of Biochemistry & Molecular Pharmacology, NYU School of Medicine, New York, NY 10016, USA; Department of Microbiology, NYU School of Medicine, New York, NY 10016, USA; Skirball Institute of Biomolecular Medicine, Departments of Cell Biology, Physiology & Neuroscience and Psychiatry, NYU School of Medicine, New York, NY 10016, USA

## Abstract

The mTOR pathway integrates both extracellular and intracellular signals and serves as a central regulator of cell metabolism, growth, survival and stress responses. Neurotropic viruses, such as herpes simplex virus-1 (HSV-1), also rely on cellular AKT-mTORC1 signaling to achieve viral latency. Here, we define a novel genotoxic response whereby spatially separated signals initiated by extracellular neurotrophic factors and nuclear DNA damage are integrated by the AKT-mTORC1 pathway. We demonstrate that endogenous DNA double-strand breaks (DSBs) mediated by Topoisomerase 2β-DNA cleavage complex (TOP2βcc) intermediates are required to achieve AKT-mTORC1 signaling and maintain HSV-1 latency in neurons. Suppression of host DNA repair pathways that remove TOP2βcc trigger HSV-1 reactivation. Moreover, perturbation of AKT phosphorylation dynamics by downregulating the PHLPP1 phosphatase led to AKT mis-localization and disruption of DSB-induced HSV-1 reactivation. Thus, the cellular genome integrity and environmental inputs are consolidated and co-opted by a latent virus to balance lifelong infection with transmission.

## INTRODUCTION

To preserve genomic integrity and maintain homeostasis, long-lived neurons must effectively respond to both exogenous and endogenous sources of DNA damage. DNA lesions such as altered bases, abasic sites, and single- and double-strand breaks (DSBs) contribute to neurotoxicity that is often associated with aging and neurological disorders, such as Parkinson’s disease, amyotrophic lateral sclerosis and Alzheimer’s disease (Madabhushi et al., 2014; McKinnon, 2013). Mammalian cells have evolved multiple DNA repair pathways to deal with various types of DNA damage (Hoeijmakers, 2001). For example, two major pathways are involved in DSB repair: homologous recombination (HR) and non-homologous end-joining (NHEJ). The HR repair pathway requires a sister chromatid to act as a template for faithful repair and is confined to late S and G2 phases of the cell cycle. HR takes place in conjunction with DNA replication and is not known to occur in terminally differentiated post-mitotic cells. In contrast, NHEJ is more error-prone but is active throughout the cell cycle, especially in G0/G1 and early S phases. As such, DNA DSBs cannot be accurately repaired in non-dividing, mature neurons, presumably due to the reliance on NHEJ as the predominant repair mechanism. Consequently, neurons are generally believed to gradually accumulate unrepaired DNA damage that over time, could potentially lead to the development of neurodegenerative disorders (Madabhushi et al., 2014; Suberbielle et al., 2013). Emerging evidence further suggests that the impact of DNA damage and repair is not restricted to cellular stress and brain disorders, but also influences the normal physiological processes of neurons. A recent study showed that neuronal activity triggers DNA DSB formation on promoters of multiple neuronal early response genes in a topoisomerase 2β (TOP2β)-dependent manner (Madabhushi et al., 2015). The generation of TOP2β-DNA cleavage complex (TOP2βcc) intermediates is required to activate early response genes in neurons. However, the underlying molecular circuitry regarding how neurons respond to DSBs and how DSBs impact neuronal cell biology, including responses to physiological stress like viral infection, are not fully understood.

Peripheral neurons in humans often harbor latent infections with neurotrophic alpha-herpesviruses such as herpes simplex virus (HSV) and as a result maintain episomal viral DNA genomes within their nucleus. HSV-1 and HSV-2 are highly prevalent worldwide, contributing to recurring ulcerative blisters at the mucosal surfaces at sites of infection, ocular lesions such as stromal keratitis that can lead to blindness, and in rare cases, encephalitis in newborns (Thellman and Triezenberg, 2017). Recent models estimate that in 2012, about 3.7 billion people aged 0-49 years are infected with HSV-1 (Looker et al., 2015). The latent state of the virus is classically defined as the absence of infectious virus production despite the presence of viral genomes in the neuronal nuclei. Expression of the more than 80 ORFs encoded by HSV-1 is largely suppressed through heterochromatin formation and other epigenetic mechanisms (Knipe and Cliffe, 2008). Periodically, the virus changes its relationship with the host cell, switching from a latent to a reactivated state; this results in the coordinate expression of productive cycle (lytic) genes and new synthesis of infectious virus that travels back along neuronal axons to the epithelium to undergo viral shedding (Wilson and Mohr, 2012). The ability of HSV-1 to establish and maintain a latent infection in peripheral neurons is essential for lifelong persistence and to function as a human pathogen (Wilson and Mohr, 2012). While a variety of conditions reportedly promote reactivation, including exposure to UV light, stress, fever, anxiety, and nerve trauma, it is unclear if neuronal responses to these stimuli operate independently or are integrated via a common molecular mechanism (Glaser and Kiecolt-Glaser, 2005; Warren et al., 1940; Wheeler, 1975).

Studies in small animal models, isolated ganglia and cultured neuron models have established that nerve growth factor (NGF) trophic support plays a key role in regulating latency and reactivation (Wilcox and Johnson, 1987, 1988; Wilcox et al., 1990). Significantly, continuous NGF-dependent signaling through the TrkA receptor and downstream activation of the AKT-mTORC1 signaling pathway is required to maintain HSV-1 latency (suppression of HSV-1 lytic gene expression) in isolated ganglia and in cultured neurons (Camarena et al., 2010; Cliffe et al., 2015; Du et al., 2015; Kobayashi et al., 2012b). Disruption of this signaling pathway by small molecule chemical inhibitors or by shRNA-mediated gene silencing, results in efficient HSV-1 reactivation. These studies establish that continuous growth factor signaling via the AKT-mTORC1 pathway is a critical parameter regulating the duration of viral latency in neurons.

Here, we provide evidence that exposure of latently-infected neurons to exogenous DNA damaging agents (DSB inducers) triggers HSV-1 reactivation. In contrast, signaling from low-level TOP2β-dependent endogenous DNA breaks provides a trophic factor-dependent signal to help maintain HSV-1 latency. Furthermore, inhibiting the Mre11-Rad50-NBS1 (MRN) complex or individual NHEJ factors responsible for repairing TOP2β-dependent DNA breaks (Hoa et al., 2016) stimulates HSV-1 reactivation. Significantly, both acute and low-level DNA damage converge onto the AKT-mTORC1 signaling axis to differentially regulate AKT (AKT1) phosphorylation and subcellular localization. We show that depletion of PHLPP1, the AKT Ser473-specific phosphatase (Gao et al., 2005), leads to prolonged AKT phosphorylation and nuclear mis-localization in response to DNA damage that, in turn, restricts HSV-1 reactivation. Our study provides new insight into the physiological role of AKT signaling and indicates how both neurotrophic growth factor and nuclear DNA damage signaling converge upon AKT to determine HSV-1 lifecycle decisions within host neurons.

## RESULTS

### Double-strand break (DSB) inducers cause HSV-1 reactivation in neuronal cells

To assess how HSV-1 latency is regulated in neurons requires a cell culture model that utilizes a homogenous neuronal population that faithfully recapitulates the hallmarks of latency and reactivation. Sympathetic neurons from dissociated superior cervical ganglia (SCG) neurons from E21 rat embryos can be cultured as a pure population of cells that depend upon the trophic factor NGF (Glebova and Ginty, 2005). To determine whether genotoxic stress can affect HSV-1 latency in SCG neurons, we utilized a primary neuronal culture model system for establishing HSV-1 latency *in vitro* and employed a wildtype HSV-1 strain expressing the enhanced Green Fluorescent Protein (GFP) fused to the Us11 true-late (γ2) protein (GFP-Us11) that can be detected during reactivation in living neurons (Benboudjema et al., 2003; Camarena et al., 2010; Kobayashi et al., 2012a; Linderman et al., 2017; Pourchet et al., 2017) (Figure 1A). Treatment of HSV-1 latently-infected neurons with either Bleomycin (DSB inducer, radiomimetic) or Etoposide (DSB inducer, TOP2 inhibitor) caused viral reactivation (productive viral replication leading to GFP-positive wells) at levels comparable to treatment with LY294002 (PI 3-kinase inhibitor, positive control). (Figure 1B). We confirmed that the accumulation of GFP-Us11 was detected in neurons (β3-tubulin, neuronal marker), and in cells experiencing DNA damage (γH2AX, DSB marker) by immunostaining (Figure 1C). Importantly, both DSB inducers resulted in accumulation of ICP27 viral mRNA, which encodes an essential immediate-early (IE) regulatory protein required for productive replication (Kim et al., 2012), and infectious virus production (Figures 1D and 1E). The DSB-induced HSV-1 reactivation was not due to the cells undergoing apoptosis; treatment with a pan-Caspase inhibitor, z-VAD-fmk, in the presence of Etoposide did not reduce HSV-1 reactivation levels (Figure S1A). Thus, exogenous DNA damage caused by DSB inducers can stimulate HSV-1 reactivation in latently-infected neurons.

**Figure 1.**
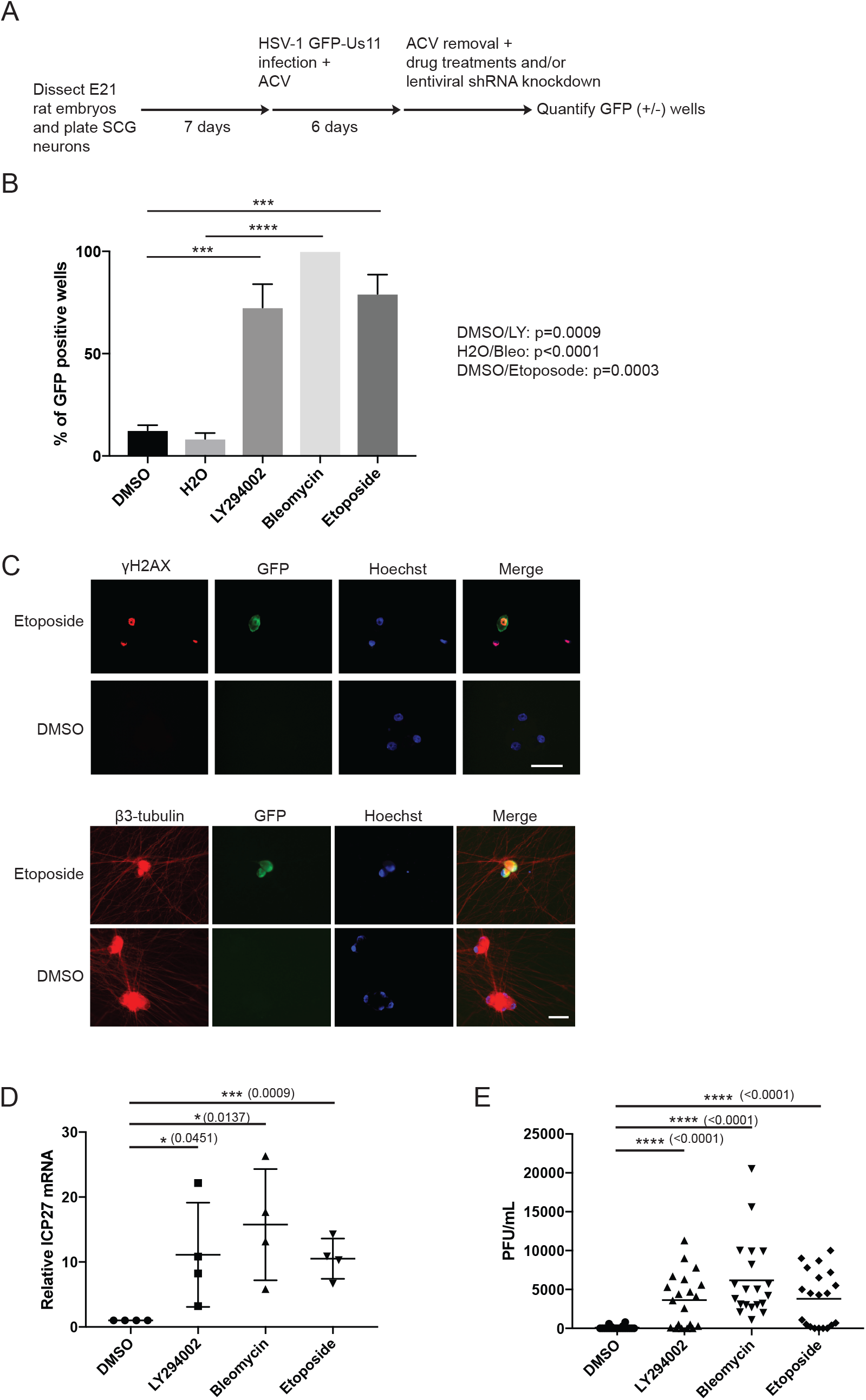
DSB inducers cause HSV-1 reactivation in primary neurons. (A) Schematic for the establishment of HSV-1 latency and reactivaton in infected rat SCG-derived neuron cultures. Dissociated SCG were seeded in 96-well plates in media supplemented with 50 ng/ml NGF for 7 days. Cells were pre-treated with 50 μM acyclovir (ACV) for 1 day prior to infection on day 8 with HSV-1 GFP-Us11. Infected cultures were maintained with ACV for an additional week to allow the virus to establish a non-replicating infection. ACV was then removed and infected cultures treated with indicated drugs or lentiviral-delivered shRNA knockdown. GFP fluorescence was monitored in live cells and quantified by percent of GFP-positive wells. 30 independently-infected wells were analyzed for individual treatments. (B) Reactivation assay comparing the response of HSV-1 GFP-Us11 infected SCG neurons treated with either Bleomycin (10 μM, 8hrs), Etoposide (10 μM, 8hrs), LY294002 (20 μM, 20hrs), or DMSO, then drug was washed out and the neurons recovered for 6 days. Data for percentage of GFP+ wells are plotted from four independent experiments with mean ±s.e.m. and p-values calculated as indicated in Experimental Procedures. (C) HSV-1 latently-infected SCG neurons were established and treated with Etoposide as in (B) for 8hrs, then drug was washed out and the neurons recovered for an additional 40hrs. Neurons were then fixed and probed with the indicated antibodies for indirect immunofluorescence (IF). Note that γH2AX levels remain present even in neurons that are GFP-negative (after Etoposide is washed out and recovered), suggesting that DNA repair is delayed in neurons and that the γH2AX signal is independent of viral replication. Nuclear DNA was visualized using Hoechst stain (blue). DMSO represents mock treatment. Bar = 50μm. (D) Latently-infected neurons were treated with the indicated chemical inducers as in (B) but were recovered for additional 12 hours, except for LY294002 treatment, and viral ICP27 mRNA levels were assessed by quantitative RT-PCR (qRT-PCR). Each data point signifies a biological replicate with mean ±s.e.m. for 4 biological replicates. (E) Infectious virus production was evaluated by plaque assay after 3 days of treatment as in (B). Bar graph shows the average number of plaque forming units (PFU) per well with mean ±s.e.m.

### MRN complex and NHEJ factors are required for HSV-1 latency

Next, the mechanism(s) by which DSB inducers cause HSV-1 reactivation was investigated. Surprisingly, we found that treatment of neuronal cells with chemical inhibitors of either the DNA-dependent protein kinase (DNA-PK) activity (NU4771), or the Mre11 nuclease activity of the MRN complex (Mirin), triggered HSV-1 reactivation in the absence of exogenous DSB inducers (Figure 2A). Both NU7441 and Mirin treatment increased the percentage of GFP-positive wells (marker of viral late gene expression), increased ICP27 viral mRNA levels, and stimulated infectious virus production (Figures 2A-C) at levels comparable to the PI 3-kinase inhibitor LY294002. Moreover, lentiviral-transduced shRNA knockdown of Ku80 (DSB-binding regulatory subunit of DNA-PK) or Rad50 (subunit of the MRN complex) using two different shRNA sequences led to HSV-1 reactivation (Figures 2D and 2E), recapitulating and reinforcing the results obtained using chemical inhibitors. Inhibition of DNA repair using chemical inhibitors or by knockdown of DNA repair factors led to an increase in γH2AX levels in the neurons, suggesting that persistent DNA damage is occurring in these cells (Figure S1B). Because we showed that both Mre11-dependent nuclease and DNA-PK activity are required for the maintenance of HSV-1 latency, it was unclear whether this involved the classical (Ku/DNA-PK) or alternative (Alt) NHEJ pathway, as the MRN complex has been implicated in the Alt-NHEJ pathway (microhomology-mediated end joining, MMEJ) (Chang et al., 2017). One major difference between the two NHEJ pathways is the DNA ligase used in the final sealing step, with the classical NHEJ requiring DNA Ligase 4 (LIG4) and Alt-NHEJ requiring DNA Ligase 3 (LIG3) (Mateos-Gomez et al., 2015; Simsek et al., 2011). To distinguish between these two possibilities, we showed that the knockdown of LIG4 using two different shRNAs caused HSV-1 reactivation in neurons, while there was no detectable effect of LIG3 knockdown (Figures 2F and 2G). Our results strongly suggest that classical NHEJ, but not Alt-NHEJ, is the critical regulator of HSV-1 latency in neurons.

**Figure 2.**
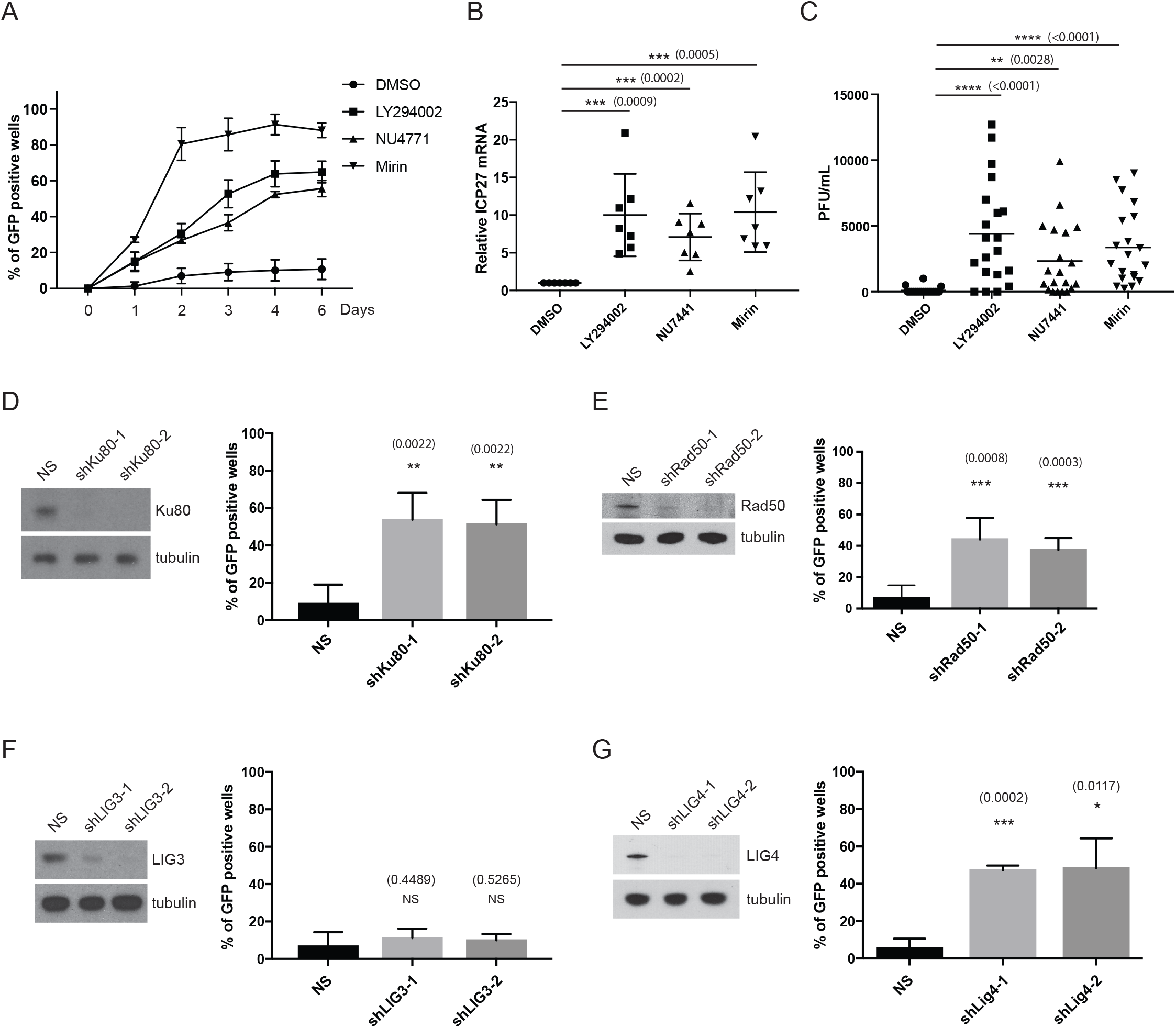
Inhibition of NHEJ or the MRN complex causes HSV-1 reactivation. (A) Reactivation assay comparing the response of HSV-1 GFP-Us11 infected SCG neurons treated with either NU4771 (1 μM, 20hrs), Mirin (100 μM, 20hrs), LY294002 (20 μM, 20hrs) or DMSO control, drugs were then washed out and the neurons were maintained for 6 days. Plot shows percentage of GFP+ wells were plotted from six independent experiments scored on successive days with mean ±s.e.m. (B) Latently-infected neurons were treated with the indicated chemicals as in (A) and ICP27 viral mRNA levels were quantified by qRT-PCR after 20 hrs of treatment. Each data point signifies a biological replicate with mean ±s.e.m. for 7 biological replicates. (C) Infectious virus production was evaluated by plaque assay after 3 days of treatment as in (A). Bar graph shows the average number of plaque forming units (PFU) per well with mean ±s.e.m. (D) Depletion of Ku80 using lentiviral-transduced shRNAs in latently-infected SCG neurons. Following ACV removal, HSV1-GFP-Us11 latently-infected cultures were infected with two different lentiviruses expressing shRNAs against rat Ku80. Cultures infected with a lentivirus expressing a non-silencing shRNA (NS) were used as a negative control. Number of wells expressing GFP was scored after 5 days with mean ±s.e.m. for 4 biological replicates. Knockdown efficiencies for individual shRNAs in latently-infected SCG neurons were confirmed by Western blot analysis (D-G). (E-G) Depletion of the indicated rat gene products (Rad50, LIG3, LIG4) using two independent lentivirus shRNAs for each were performed and scored for reactivation as in (D).

### NHEJ and MRN complex promote AKT-mTORC1 signaling to maintain HSV-1 latency

Our previous work demonstrated that the AKT-mTORC1 signaling axis is a critical regulator of HSV-1 latency in SCG neurons (see schematic diagram in Figure 3A) (Camarena et al., 2010; Kobayashi et al., 2012b). Although several earlier studies have shown that DNA-PK and MRN-dependent ATM activation can promote AKT phosphorylation on Ser473 in non-neuronal cell lines (Bozulic et al., 2008; Fraser et al., 2011; Wan et al., 2013), it is quite conceivable that NHEJ and the MRN complex could maintain HSV-1 latency independently of the AKT-mTORC1 signaling axis in primary neurons. Here, we showed that inhibition of NHEJ or MRN complex activity by chemical inhibitors or shRNA knockdown resulted in the reduction of baseline AKT phosphorylation on Ser473 (marker for AKT activation) (similar results were obtained for phosphorylation of AKT on Thr308) (Figures 3B and 3C, other data not shown); AKT phosphorylation on these residues is critical for AKT downstream activity, with mTORC1 kinase being a key effector (Manning and Toker, 2017). In addition, ATM inhibition using the chemical inhibitor KU55933 reduced AKT Ser473 phosphorylation and led to an increase in HSV-1 reactivation at levels comparable to Mirin treatment (Figure S3A). Interestingly, the effects of DNA repair inhibition on AKT phosphorylation appears to be independent of viral infection (Figure 3B). Other non-neuronal, dividing cells, such as primary rat embryonic fibroblasts (REFs) and an established rat fibroblast cell line (Rat2), were independently tested and were shown to have reduced baseline AKT phosphorylation when compared to rat SCG neurons, and remarkably, perturbation of the MRN complex or NHEJ factors had no detectable effect on baseline AKT phosphorylation status in these different rat cell lines (Figures S1C-E, other data not shown).

**Figure 3.**
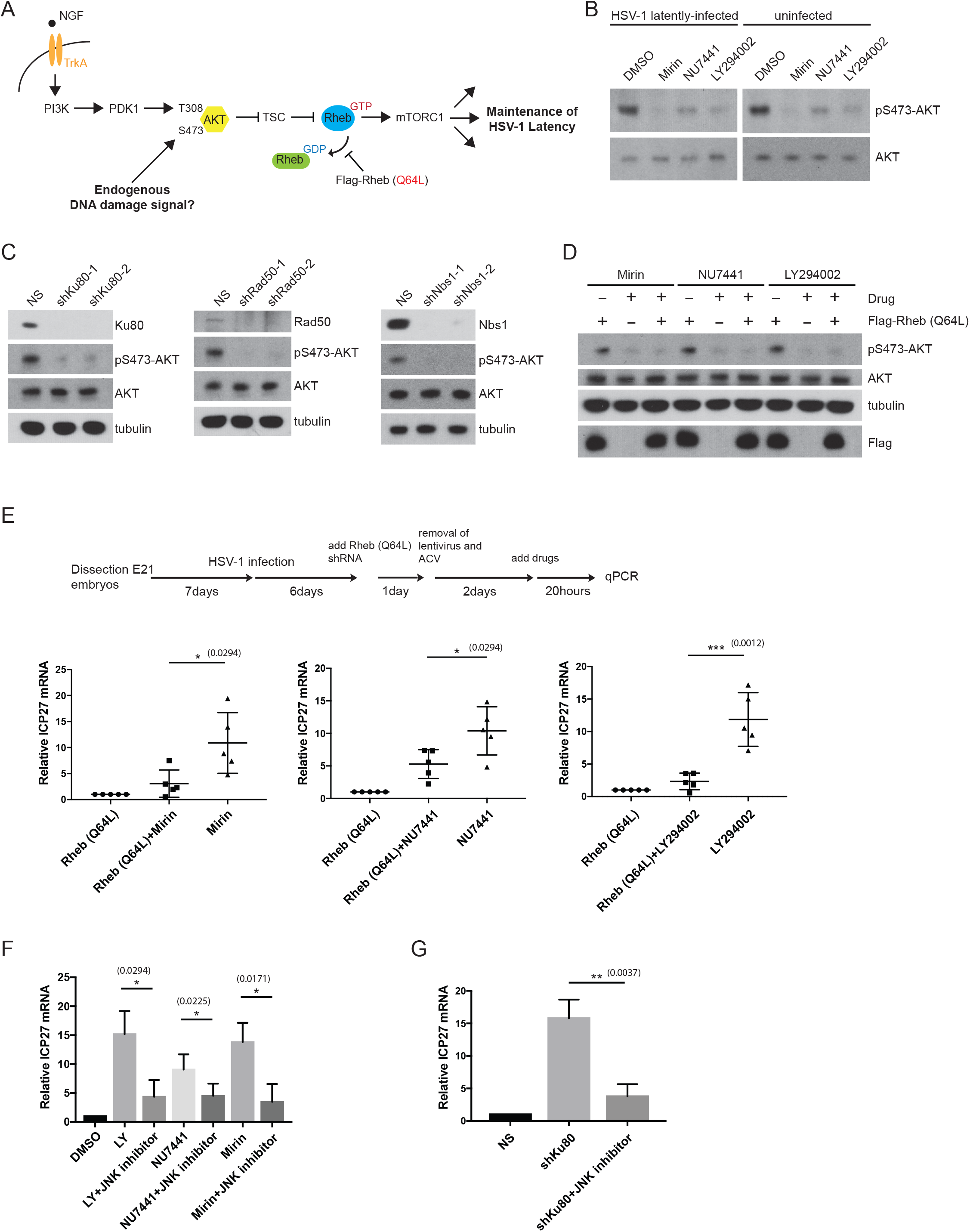
DNA repair factors maintain HSV-1 latency through the canonical AKT-mTORC1 signaling axis. (A) Schematic diagram of mTORC1 signaling induced by both the NGF-mediated TrkA/PI3K/PDK1/AKT pathway and DNA damage (and DNA repair inhibition) activation, leading to the maintenance of HSV-1 latency. (TSC) Tuberous sclerosis complex, (Rheb) Rheb is depicted in GDP- and GTP-bound forms. The Rheb (Q64L) mutant is a constitutively active protein that acts at the point shown (stabilization of the GTP-bound form) to bypass upstream AKT inhibition in order to activate the downstream mTORC1 signaling pathway. (B) Western blot analysis comparing levels of AKT Ser473 phosphorylation in uninfected and HSV-1 latently-infected SCG neurons treated with either NU4771 (1 μM, 20hrs), Mirin (100 μM, 20hrs), LY294002 (20 μM, 20hrs) or DMSO control. LY294002 serves as a positive control for inhibiting AKT Ser473 phosphorylation. (C) Different DNA repair factors are depleted by shRNAs for 5 days in uninfected SCG neurons and analyzed by Western blot for AKT Ser473 phosphorylation as in (B) and along with other indicated proteins. (D) Uninfected SCG neurons were transduced with either empty vector or Flag epitope-tagged Rheb (Q64L) mutant expression construct and treated as in (B) and analyzed by Western blot with the indicated antibodies. (E) Lentivirus-delivered expression of the constitutively active Flag-Rheb (Q64L) mutant in HSV-1 latently-infected SCG neurons in the presence or absence of treatment with Mirin, NU7441 or LY294002 for 20 hrs as indicated (see schematics). Viral ICP27 mRNA levels were then quantified by qRT-PCR. Each data point signifies a biological replicate with mean ±s.e.m. for 5 biological replicates. (F-G) Induction of ICP27 viral mRNA in response to the different DNA repair inhibitors (F) or shRNA knockdown of Ku80 (F) can be blocked by a JNK inhibitor. LY294002 serves as a positive control for the JNK inhibitor consistent with a previous study (Cliffe et al., 2015). Relative ICP27 mRNA levels were quantified by qRT-PCR after 20 hrs drug treatment and represent the mean ±s.e.m. for 4 biological replicates.

To determine whether HSV-1 reactivation upon NHEJ or MRN complex inhibition was specific for the AKT-mTORC1 signaling pathway, but not other AKT effector pathways, we utilized a Rheb Gln64 to Leu mutant (Q64L) that locks the the Rheb GTPase into a GTP-bound state; the Rheb Q64L mutant behaves as a constitutive mTORC1 activator (Martz et al., 2014). Thus, the Rheb Q64L bypasses the requirement for AKT phosphorylation and/or its upstream regulators (see schematics in Figure 3A). Our previous work demonstrated that Rheb Q64L mutant expression is sufficient to inhibit LY294002-mediated HSV-1 reactivation (Kobayashi et al., 2012b). Here, we show that Rheb Q64L mutant expression also reduced HSV-1 reactivation in response to NHEJ or MRN inhibition, overcoming both Mirin and NU7441-induced AKT inhibition in neurons (Figures 3D and 3E).

Moreover, we found that HSV-1 ICP27 mRNA accumulation triggered by inhibiting DNA repair (Mirin or NU7441 treatment or Ku80 shRNA) was sensitive to treatment with a Jun N-terminal kinase (JNK) inhibitor, JNK Inhibitor II (Figures 3F and 3G). This is significant because the initiation of HSV-1 gene expression through the inhibition of PI 3-kinase during reactivation was shown to be dependent upon a methyl/phospho switch on viral lytic gene promoters that requires JNK activity (Cliffe et al., 2015). Collectively, these data establish that both NHEJ and the MRN complex activity are required for the maintenance of HSV-1 latency; they act through the canonical AKT-mTORC1 pathway to prevent JNK signaling for the repression of HSV-1 lytic genes.

### TOP2β and TDP2 is required for the maintenance of HSV-1 latency

Although both the MRN complex and NHEJ factors are required to maintain HSV-1 latency, the source of endogenous DNA damage that signals to the AKT-mTORC1 pathway remains unclear. TOP2 enzymes resolve DNA catenanes by catalyzing the transient formation of a DSB, enabling an intact DNA duplex to pass through the DSB, followed by a re-ligation (Nitiss, 2009a). During such transient DSB formation, TOP2 becomes covalently bound to the 5’ DNA end of the break, forming TOP2cc intermediates. Failure to complete the re-ligation step and release of TOP2 produces a DSB. The protein-DNA covalent complex associated with TOP2cc lesions can be removed by either tyrosyl-DNA-phosphodiesterase 2 (TDP2) or the MRN complex, followed by the repair of the DSB by the NHEJ pathway (Aparicio et al., 2016; Cortes Ledesma et al., 2009; Gao et al., 2014; Hoa et al., 2016; Zeng et al., 2011). The generation and stabilization of TOP2cc-DSBs is an important aspect of cancer therapy (Nitiss, 2009b; Pommier, 2013). Nearly half of the current antitumor regimens consist of treatment with topoisomerase inhibitors, including Etoposide, a TOP2 poison which strongly stabilizes TOP2cc intermediates and leads to the formation of stable DSBs. Previous studies have shown that Mre11 contributes to the canonical NHEJ-dependent repair of Etoposide-induced DSBs, and that Ku proteins and Mre11 functions epistatically in this pathway (Aparicio et al., 2016; Hoa et al., 2016). Moreover, TOP2β was identified as a critical regulator of neuronal activity-induced DSBs at promoters of early-response genes and necessary for the expression of genes involved in neuronal growth (Madabhushi et al., 2015).

To test whether TOP2β or TDP2 plays a role in regulating HSV-1 reactivation, we used two different shRNAs to knockdown endogenous TOP2β or TDP2 in SCG neurons. The depletion of either TOP2β or TDP2 led to a decrease in baseline AKT Ser473 phosphorylation (Figures 4A and 4D). The loss of TOP2β also led to a decrease in γH2AX signal (Figure 4A), this suggests that the DSBs generated by TOP2βcc intermediates represent the major source of endogenous DNA damage required for continuous AKT activation. Indeed, we found that inhibiting Mre11-dependent nuclease activity by Mirin treatment caused an increase in the abundance of TOP2βcc in SCG neurons (Figure 4B). As a control, knockdown of TOP2β reduced TOP2βcc intermediates both in SCG neurons and non-neuronal Rat2 fibroblasts (Figures 4B and S2A). Consistent with this hypothesis, TOP2β is required for maintaining HSV-1 latency as TOP2β depletion by shRNAs in latently-infected SCG neurons resulted in accumulation of viral lytic mRNA markers of reactivation (Figure 4C). Under our conditions, TOP2β depletion does not affect IE viral gene expression for reactivation and during the acute infection (lytic) cycle, suggesting that IE viral gene expression program can occur independently of TOP2β (Figures 4C and S2C). However, other viral maturation processes involved in HSV-1 reactivation, such as expression of viral late genes (as exemplified by GFP-Us11 accumulation) were not observed in TOP2β-depleted neurons (Figure S2B); this could be due to a critical role for TOP2β in relaxing supercoiled viral DNA during viral DNA replication. Nevertheless, the loss of TDP2 led to an increase in HSV-1 reactivation (GFP-Us11 accumulation) (Figure 4D). In contrast to TOP2β, TDP2 is not necessary for HSV-1 lytic replication.

**Figure 4.**
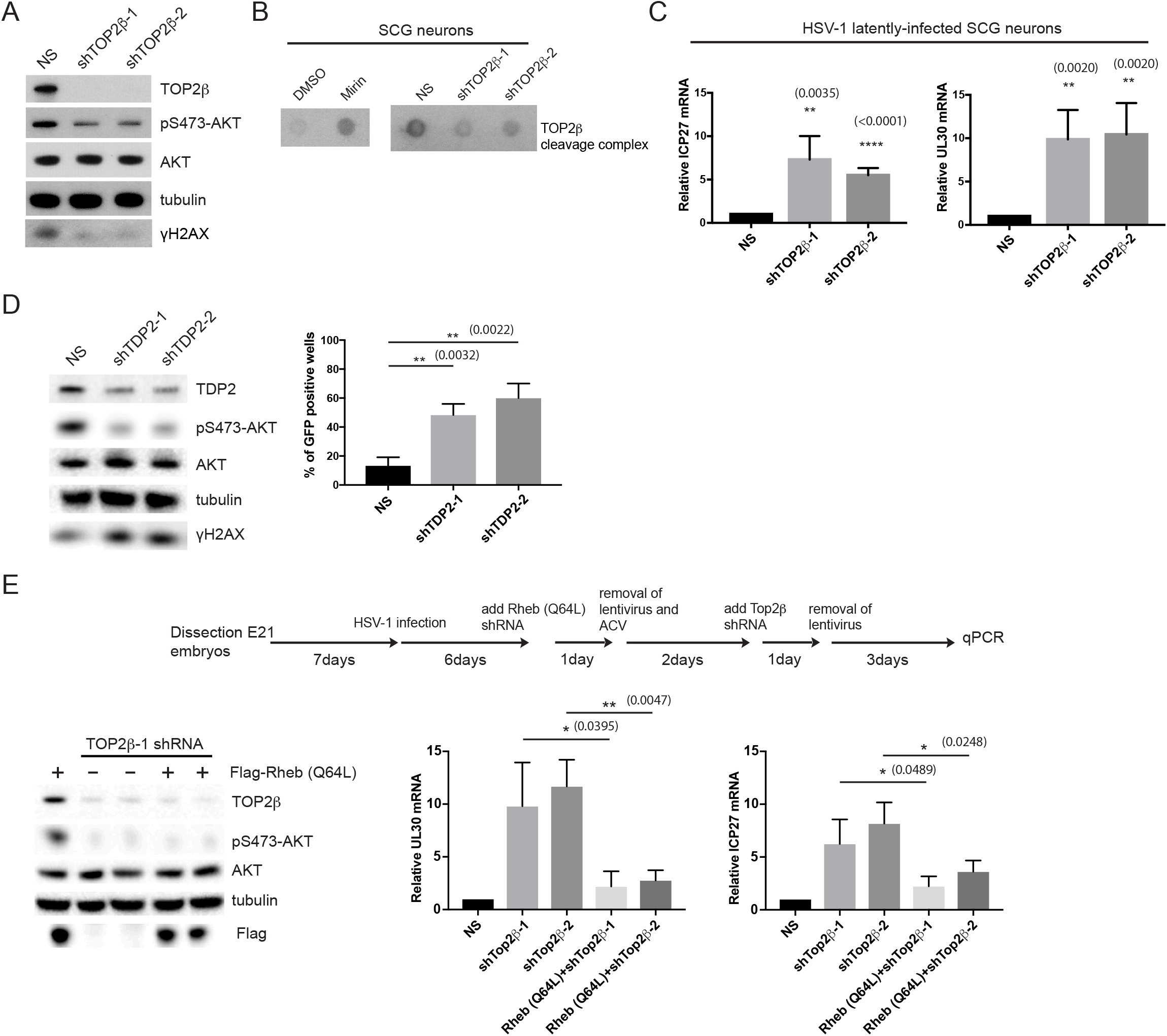
Generation of endogenous TOP2βcc intermediates is critical for the maintenance of HSV-1 latency. (A) Uninfected SCG neurons were transduced with two different shRNAs against TOP2β and analyzed by Western blot for differences in AKT and H2AX phosphorylation levels. (B) TOP2β covalently-bound to genomic DNA from uninfected SCG neurons were isolated and displayed using a Dot blot method. Cells were treated either with Mirin or depleted of TOP2β with the indicated shRNAs. (C) HSV-1 latently-infected SCG neurons were transduced with non-silencing (NS) or TOP2β shRNAs for 3 days and scored for ICP27 or UL30 viral mRNA levels by qRT-PCR with the mean ±s.e.m. for 4 biological replicates. (D) Uninfected SCG neurons were transduced with two different shRNAs against TDP2 and analyzed by Western blot for differences in AKT and H2AX phosphorylation levels (left panel). HSV-1 latently-infected SCG neurons transduced with either NS or two different TDP2 shRNAs were scored for GFP-positive neurons after 5 days (right panel). (E) Uninfected SCG neurons were transduced with Flag-Rheb (Q64L) and then shRNA against TOP2β to monitor relative efficiency of knockdown of TOP2β and Flag-Rheb (Q64L) expression by Western blot analysis (left panel). HSV-1 latently-infected SCG neurons were transduced with either empty vector or Flag-Rheb (Q64L), and then with two different shRNAs against TOP2β (see schematics) and scored for UL30 or ICP27 viral mRNA levels by qRT-PCR with the mean ±s.e.m. for 4 biological replicates (right panels).

Based on the reduced AKT Ser473 phosphorylation upon TOP2β knockdown (Figure 4A), it is highly suggestive that the loss of TOP2βcc intermediates leads to HSV-1 reactivation through the inhibition of the canonical AKT-mTORC1 signaling pathway. To demonstrate this experimentally, we asked whether the expression of the Rheb Q64L mutant in TOP2β-deficient neurons could overcome AKT inhibition and suppress HSV-1 reactivation. Indeed, Rheb Q64L expression in TOP2β knockdown cells reduced expression of both the UL30 and ICP27 viral transcripts (Figure 4E). Thus, our data supports a model whereby both TOP2β and TDP2 are major regulators of HSV-1 latency in neurons via the generation of TOP2βcc intermediates, which is somehow required for the continuous DDR-mediated activation of the AKT-mTORC1 pathway.

### Acute DSB inducers are unable to promote sustained AKT activation in neuronal cells

In neurons, NGF-dependent activation of the TrkA receptor stimulates PI 3-kinase activity, which leads to downstream phosphorylation of two key residues on AKT1, Thr308 in the T-loop of the catalytic core, and Ser473 in the C-terminal hydrophobic motif (Camarena et al., 2010; Manning and Toker, 2017). The phosphoinositide-dependent protein kinase 1 (PDK1) phosphorylates AKT1 on Thr308, which is required for AKT activity, but maximal activation of AKT also requires phosphorylation of Ser473 (Manning and Toker, 2017). The major AKT Ser473 kinase is the mTOR complex 2 (mTORC2). However, other kinases can perform this function including DNA-PK and ATM, both members of the PI 3-kinase family, which phosphorylates AKT Ser473 in response to DNA damage (Bozulic et al., 2008; Fraser et al., 2011). In agreement with an early study (Bozulic et al., 2008), we found that the treatment of uninfected SCG neurons with either NU7441, or alternatively by Ku80 shRNA knockdown, inhibited Bleomycin-or Etoposide-induced AKT Ser473 phosphorylation (Figures 5A and 5B). Despite the dramatic loss of DNA damage-induced AKT phosphorylation in the absence of DNA-PK activity, the DNA damage signal (γH2AX) remained elevated, which suggests that AKT activation can be uncoupled to the sensing of DSBs via H2AX phosphorylation (Figure 5B). In contrast, inhibiting ATM had minimal effect on Etoposide-induced AKT Ser473 phosphorylation (Figure S3B), even though ATM has been reported to mediate the repair of DSBs with blocked DNA ends (Álvarez-Quilón et al., 2014). Interestingly, inhibition of RICTOR, a subunit of mTORC2, caused a reduction in baseline AKT Ser473 phosphorylation in untreated neurons, but had no detectable effect on DNA damage-induced AKT Ser473 phosphorylation (Figure 5C). In line with this result, loss of RICTOR caused spontaneous HSV-1 reactivation in latently-infected neurons (Figure S4A). This argues strongly that mTORC2 participates in the basal activation of AKT in response to neurotrophic growth factor signaling, which influences HSV-1 latency.

**Figure 5.**
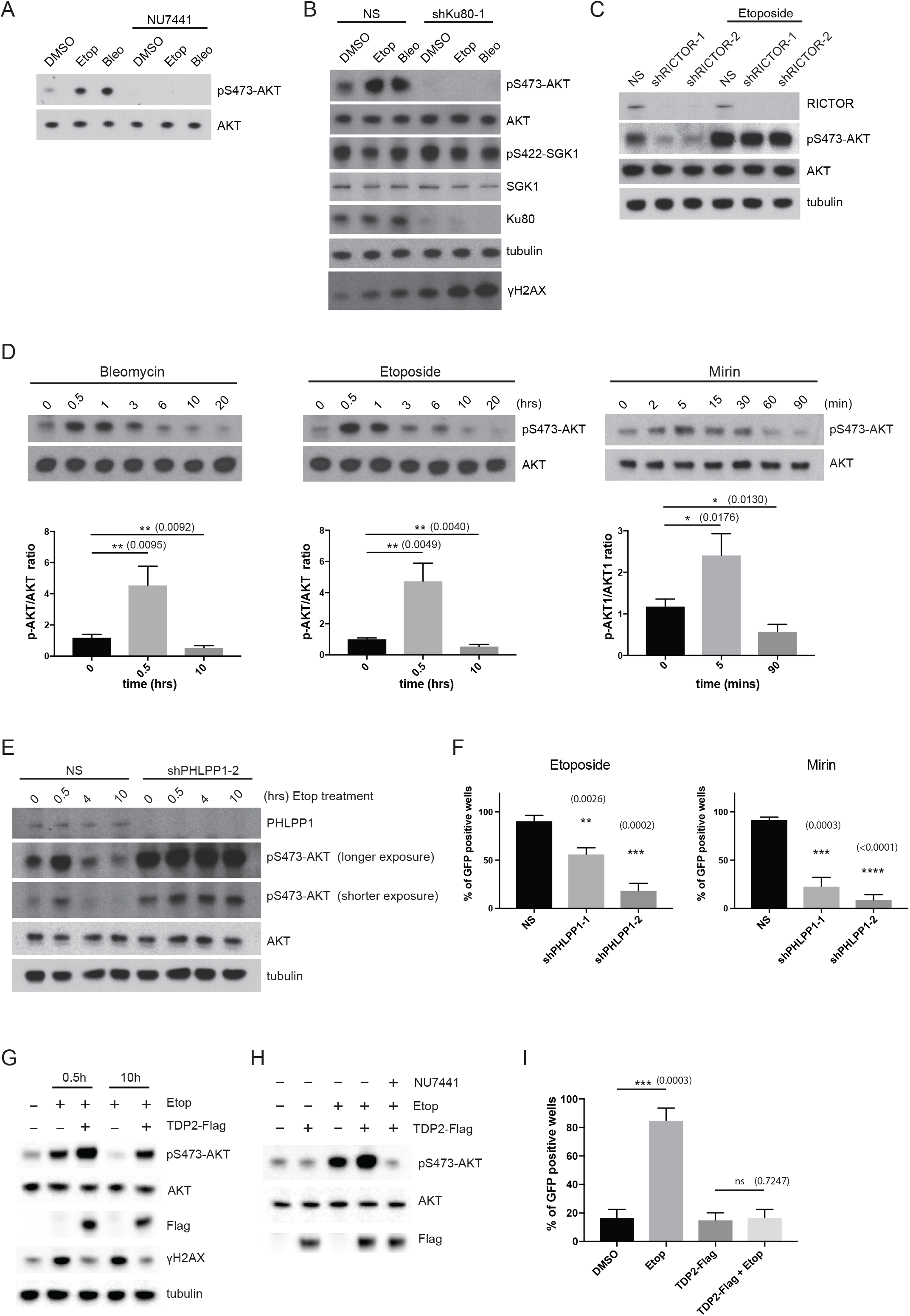
Loss of PHLPP1 causes sustained AKT Ser473 phosphorylation and reduced HSV-1 reactivation in response to DSB inducers. (A) Treatment of uninfected SCG neurons with NU7441 (DNA-PK activity inhibitor) prevents basal and DSB-induced AKT Ser473 phosphorylation. Samples were analyzed by Western blot with the indicated antibodies. (B) Uninfected SCG neurons were transduced with a lentivirus expressing a non-silencing shRNA (NS) or an shRNA specific for Ku80. Following treatment with either DMSO (control), Etoposide, or Bleomycin, total protein was analyzed by immunoblotting using the indicated antibodies. Phosphorylation of SGK1 (another substrate of AKT) served as a negative control. (C) Knockdown of RICTOR, a subunit of the mTORC2 complex, using two different lentivirus-delivered shRNAs had no detectable effect on Etoposide-induced AKT Ser473 phosphorylation. Samples were analyzed by Western blot with the indicated antibodies. (D) Time-course study performed using DSB inducers (Bleomycin, Etoposide) or the DNA repair inhibitor (Mirin) to analyze the kinetics of AKT Ser473 phosphorylation. Samples were analyzed by Western blot (top panels, a representative image of pS473-AKT and AKT signals) and quantified by ImageJ for the corresponding DSB inducers at the indicated time-points (bottom panels). The signal was normalized to endogenous AKT levels (pS473-AKT/AKT) as a ratio, with mean ±s.e.m. for 3 biological replicates. (E) PHLPP1 depletion by lentivirus-delivered shRNA causes elevated and sustained AKT phosphorylation. Samples were analyzed by Western blot and probed with indicated antibodies. (F) Both Etoposide- and Mirin-induced HSV-1 reactivation are inhibited by PHLPP1 depletion using two different shRNAs. Number of GFP-positive wells was scored after 6 days with mean ±s.e.m. for 3 biological replicates. Knockdown efficiency of PHLPP1 in latently-infected SCG neurons were confirmed by Western blot analysis (see Figure S4C). (G) TDP2-Flag overexpression hyper-activates AKT Ser473 phosphorylation in response to Etoposide treatment. Uninfected SCG neurons were transduced with lentivirus expressing Flag epitope-tagged TDP2 (TDP2-Flag) for 3 days and then treated with Etoposide for the indicated times and analyzed by Western blot. (H) TDP2-dependent hyper-activation of AKT Ser473 phosphorylation by Etoposide requires DNA-PK activity. Uninfected SCG neurons were transduced with TDP2-Flag for 3 days and then treated with Etoposide in the presence or absence of NU7441 as indicated and analyzed by Western blot. (I) TDP2-Flag overexpression inhibits Etoposide-induced HSV-1 reactivation. HSV-1 latently-infected SCG neurons were transduced with TDP2-Flag for 3 days and then treated with Etoposide for 8hrs as indicated. Number of GFP-positive wells was scored after 3 days with mean ±s.e.m. for 3 biological replicates.

Since treatment of uninfected SCG neurons with either Bleomycin or Etoposide can induce robust AKT Ser473 phosphorylation, it is paradoxical why HSV-1 reactivation would occur. Intuitively, hyper-phosphorylation of AKT should limit, but not induce, HSV-1 reactivation due to the hyper-activation of the AKT-mTORC1 signaling pathway. Unexpectedly, we found that both Bleomycin and Etoposide treatment induced only transient AKT phosphorylation over time in neurons, peaking early at 30 min to 1 hour, but after 10 hrs, the AKT phosphorylation status fell below baseline (untreated) levels (Figure 5D). In contrast, Etoposide-induced AKT phosphorylation levels remained elevated even after 10 hrs in Rat2 fibroblasts (Figure S4B). Inhibiting Mre11 nuclease activity with Mirin treatment yielded a similar trend to the dynamics of AKT phosphorylation with the acute DSB inducers, namely a rapid increase in AKT phosphorylation (peak at 5 min), followed by a decrease in signal below baseline level after 90 min (Figure 5D). Thus, our data supports the findings that acute DSB inducers (and DNA repair inhibitors) can cause HSV-1 reactivation in neurons due to a failure to sustain elevated levels of AKT phosphorylation.

The inability of DSB inducers to sustain elevated AKT activation in SCG neurons could be due in part to differential regulation by an AKT phosphatase. The PH domain leucine-rich repeat protein phosphatase, PHLPP (PHLPP1), is reported to specifically dephosphorylates AKT1 Ser473 (Gao et al., 2005). We therefore tested whether PHLPP1 knockdown using shRNAs was sufficient to modulate AKT phosphorylation in neurons. Strikingly, loss of PHLPP1 led to an elevated and sustained AKT phosphorylation state (at least up to 10 hrs) in response to Etoposide treatment (Figures 5E and S4C). A second family member, PHLPP2, which has also been implicated as an AKT phosphatase (Brognard et al., 2007), had little or no effect on AKT phosphorylation status in this context (Figure S4D). Significantly, depletion of PHLPP1 inhibited HSV-1 reactivation by either Etoposide or Mirin treatment, demonstrating the relevance of AKT phosphorylation dynamics in the regulation of the HSV-1 latent-lytic switch. Notably, even though PHLPP1 depletion caused a slight increase in steady-state AKT phosphorylation levels (Figure S4C), this had no measurable impact on HSV-1 latency or lytic replication (Figures S4E-G). Based on these observations, we identify PHLPP1 as a new regulator of HSV-1 reactivation in neurons.

It has been previously shown that overexpressing TDP2 can reduce Etoposide-mediated TOP2cc intermediates and chromosomal aberrations (Hoa et al., 2016). How TDP2 regulates AKT-mTORC1 signaling remains elusive. Interestingly, we found that overexpressing TDP2 accentuated AKT Ser473 phosphorylation in the presence of Etoposide (Figure 5G). This increase in phosphorylation level was dependent on DNA-PK activity as treatment with NU7441 can reduce AKT Ser473 phosphorylation (in the presence of Etoposide and overexpressed TDP2) back to the baseline level (Figure 5H). Even though TDP2 overexpression dramatically elevated AKT Ser473 phosphorylation, it was still susceptible to downregulation after 10 hrs of Etoposide treatment (Figure 5G). Nevertheless, at 10 hrs while in the presence of Etoposide and overexpressed TDP2, the level of AKT Ser473 phosphorylation was still significantly higher than baseline untreated levels (Figure 5G). In line with this observation, TDP2 overexpression was capable of inhibiting Etoposide-induced HSV-1 reactivation in SCG neurons (Figure 5I). How TDP2 stimulates DNA-PK activity to activate AKT in this context is completely unknown. It is noted that TDP2 overexpression in the presence of Etoposide led to a decrease in γH2AX signal in comparison to Etoposide treatment alone (Figure 5G), suggesting that TOP2βcc intermediates are being processed and repaired while AKT is being activated by DNA-PK.

### NGF-dependent cytoplasmic signaling is required for transient DSB-induced AKT nuclear localization and activation

AKT is a well-established central hub for cellular signal transduction, relaying critical information generated at the cell surface to nuclear functions (Manning and Toker, 2017). This process relies mainly on a series of phosphorylation events: cell surface receptor activation by extracellular growth factor binding, activation of PI 3-kinase, and phosphorylation of AKT on Thr308 by PDK1 and Ser473 by mTORC2. Activated AKT then phosphorylates several downstream signaling molecules, including tuberous sclerosis complex 2 (TSC2), the Forkhead Box O (FOXO) transcription factors, Glycogen Synthase Kinase 3 (GSK3) and others (Manning and Toker, 2017). In contrast, precisely how nuclear DNA damage elicits phosphorylation and activation of the AKT-mTORC1 signaling pathway in a spatial and temporal manner remains unknown. To determine whether AKT localization changes in response to DSB inducers, we treated SCG neurons with Etoposide in a time-course experiment and analyzed the subcellular distribution of AKT using immunofluorescence confocal microscopy. In untreated neurons, we found AKT to be either distributed primarily in the cytoplasm (nuclear exclusion), including axons, or evenly distributed between the nuclear/cytoplasmic compartments (Figure 6A, see 0h panels for representative images). Strikingly, after just 30 min of Etoposide treatment, most of the cells displayed a distinct nuclear accumulation (Figure 6A, 0.5h panel). After 4 hrs of Etoposide treatment, the signal became more evenly distributed throughout the soma (Figure 6A, 4h panel). Whether the neurons were latently-infected or uninfected with HSV-1 had no detectable bearing on AKT localization dynamics in response to Etoposide treatment (Figures 6A and S5A). The cytoplasmic localization of the downstream regulator and substrates of AKT, such as mTOR or TSC2, did not detectably change in response to Etoposide treatment (Figure 6A). This suggests that AKT subcellular localization is dynamically regulated as a means to receive a nuclear DNA damage signal which, in turn, can activate a cytoplasmic signaling event. Using a pre-permeabilization/extraction protocol (allows for the enrichment of chromatin-bound proteins and removal of soluble proteins in intact cells), we observed AKT in nuclear foci that co-localizes with γH2AX (a marker for DSBs) following Etoposide treatment at the 30 min time-point (Figure 6B). This is in agreement with a previous study showing that AKT can form nuclear foci in non-neuronal cell lines in response to ionizing radiation (Bozulic et al., 2008). Based on these results, we predict that the phosphorylation status of AKT could be a critical determinant on its sub-cellular localization. In support of this idea, we showed that depleting PHLPP1 (AKT Ser473-specific phosphatase) caused prolonged nuclear accumulation of AKT, even after 10 hrs of Etoposide treatment (Figure 6C). Thus, the phosphorylation status of AKT strongly correlates with its subcellular localization in response to DNA damage signals.

**Figure 6.**
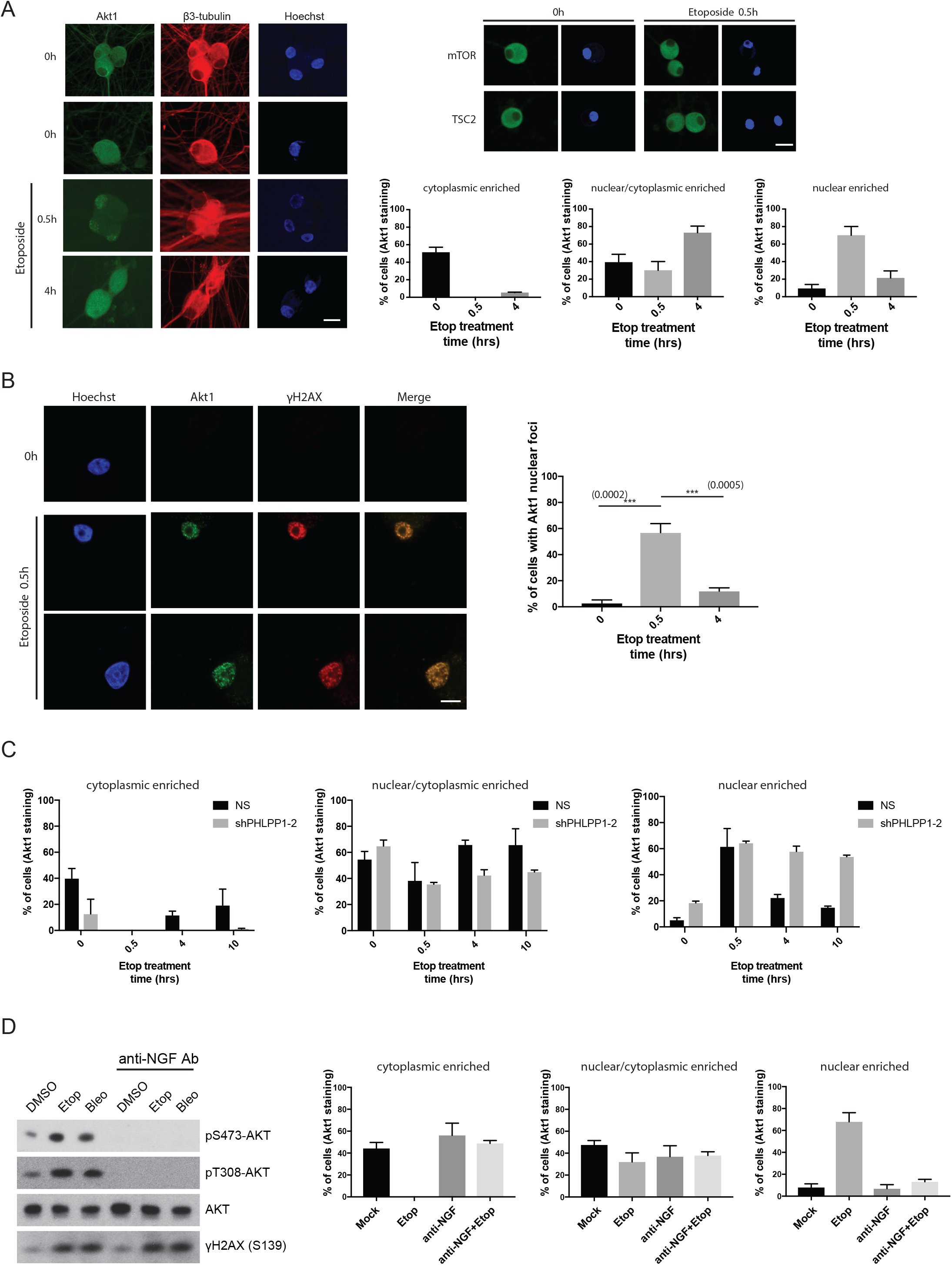
AKT subcellular localization is controlled by both extracellular growth factor and nuclear DSB signaling. (A) Confocal microscopy images of cultured, uninfected SCG neurons that were untreated (0h) or treated with Etoposide for the indicated times (h) and subsequently immuno-stained with the indicated antibodies. Nuclear DNA was visualized using Hoechst stain (blue). The percentage of cells with either cytoplasmic, nuclear/cytoplasmic, or nuclear enriched AKT1 subcellular localization were quantified and displayed as bar graphs with mean ±s.e.m. for 3 biological replicates. Representative images of AKT1 staining for cytoplasmic enriched (top panel, 0h), nuclear/cytoplasmic enriched (middle panel, 0h), and nuclear enriched (bottom panel, Etoposide 0.5h) are shown. Bar = 20μm. Images acquired from parallel cultures immuno-stained with anti-mTOR or TSC2 provide specificity controls indicating that the subcellular distribution of two downstream AKT effectors did not detectably change in response to Etoposide. (B) Uninfected SCG neurons treated with Etoposide for the indicated time (h) were processed to detect chromatin-bound proteins in intact cells. Neurons were immuno-stained with the indicated antibodies and representative images of AKT1 and γH2AX co-localization are shown. The percentage of cells displaying 5 or more AKT1 nuclear foci per nuclei are quantified with mean ±s.e.m. for 3 biological replicates. Bar = 10μm. (C) Comparing the impact of PHLPP1 shRNA knockdown *versus* non-silencing (NS) control shRNA on AKT subcellular distribution in uninfected SCG neurons treated with Etoposide for the indicated times (hrs). The percentage of cells with either cytoplasmic, nuclear/cytoplasmic, or nuclear enriched AKT1 subcellular localization were quantified and displayed as bar graphs with mean ±s.e.m. for 3 biological replicates. (D) Cultured SCG neurons were treated with or without anti-NGF blocking antibody in the media to inhibit TrkA receptor activation and then treated with DMSO (vehicle control) or DSB inducers as indicated. Samples were analyzed by Western blot to ensure that anti-NGF antibody blocked AKT phosphorylation, but not DSB formation (γH2AX). The subcellular distribution of AKT was analyzed by immuno-staining and confocal microscopy imaging. The percentage of cells with either cytoplasmic, nuclear/cytoplasmic, or nuclear enriched AKT1 subcellular localization were quantified and displayed as bar graphs with mean ±s.e.m. for 3 biological replicates.

Extracellular NGF signaling via TrkA receptor activates PI 3-kinase and PDK1-dependent phosphorylation of AKT on Thr308. This phosphorylation step is essential for priming AKT to be further phosphorylated on Thr308 and Ser473, and for signal amplification of the AKT-mTORC1 pathway (Manning and Toker, 2017). To define how extracellular NGF signaling to AKT might impact DNA damage-induced AKT activation, NGF signaling was neutralized with anti-NGF antibody in the presence (or absence) of the DSB inducer, Etoposide. Treatment of SCG neurons in the media with the anti-NGF antibody caused a dramatic loss of both steady-state (untreated) and DNA damage-induced phosphorylation of Thr308 and Ser473 on AKT (Figure 6D). Importantly, in the absence of AKT phosphorylation due to anti-NGF antibody pretreatment (inhibiting TrkA receptor), AKT could no longer accumulate in the nucleus upon Etoposide treatment (Figure 6D). This strongly suggests that extracellular signaling from neurotrophic growth factors can inform and contribute towards the cellular response to nuclear DNA damage via the AKT-mTORC1 signaling axis (model, Figure 7).

**Figure 7.**
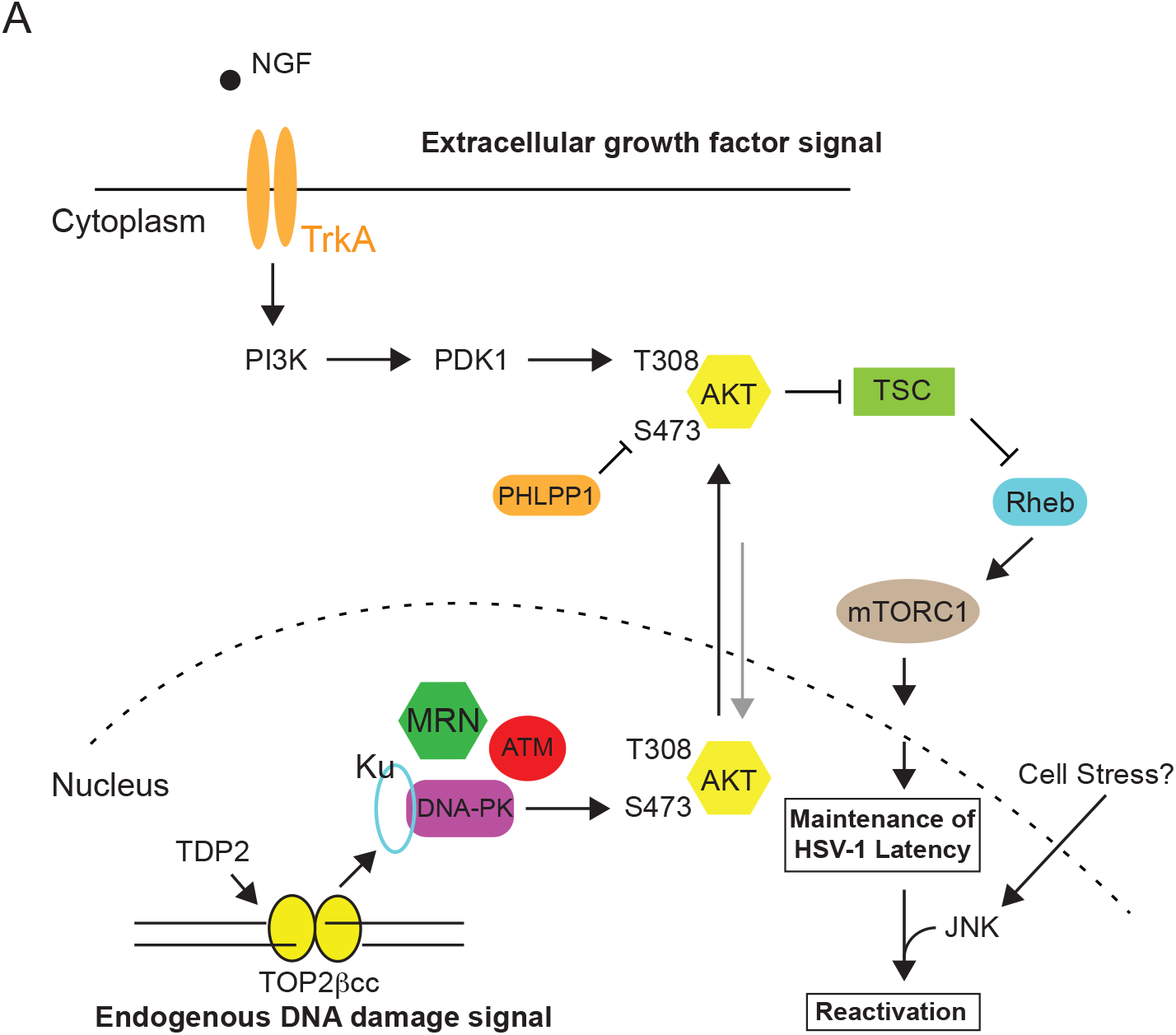
A working model of how the spatial distribution of nuclear DNA damage and extracellular neurotrophic factor signaling coordinates AKT-mTORC1 activation. (A) Working model depicting how both extracellular growth factor and TOP2β-mediated nuclear DNA damage signaling collaborate together to sustain AKT-mTORC1 activation to maintain HSV-1 latency. NGF-dependent activation of TrKA leads to sustained AKT Thr308 phosphorylation. This step is critical in order for AKT to be subsequently phosphorylated on Ser473 by DNA-PK. DNA-PK is then activated from transient DSBs generated by TOP2βcc intermediates in collaboration with TDP2, which can occur at promoters of early-response host genes during normal neuronal activity (Madabhushi et al., 2015). The MRN complex is critical for the processing of TOP2βcc for both DNA repair (Hoa et al., 2016) and the activation of DNA-PK (our study). Based on our present study, we speculate that AKT undergoes continuous nucleocytoplasmic shuttling. AKT is likely phosphorylated on Thr308 at the plasma membrane by PDK1. This step is critical for initiating AKT nuclear translocation and phosphorylation by DNA-PK on Ser473, but negatively regulated by PHLPP1, leading to its nuclear export and activation of cytoplasmic AKT-mTORC1 canonical pathway. Continuous AKT-mTORC1 signaling is crucial for the maintenance of HSV-1 latency. Thus, the sustained activation of the AKT-mTORC1 signaling axis in neurons requires consolidation from two independent signals generated from potentially different cellular compartments (nuclear and plasma membrane/cytoplasm). During latency, histones at HSV-1 lytic promoters remain in a repressed state. Neuronal cell stress stimuli triggering DLK/JIP3-mediated activation of JNK will contribute to histone demethylation and a histone methyl/phospho switch (Cliffe et al., 2015), leading to euchromatin-associated marks enriched on lytic promoters during reactivation (not pictured). The novel link between neurotrophic factors and neuronal genome integrity is an unexplored territory for neuronal homeostasis. Nevertheless, neurotropic viruses have conveniently co-opted this genome integrity link for its own life cycle advantages.

## DISCUSSION

In addition to their critical roles in balancing anabolic and catabolic responses, AKT and its downstream effector mTORC1 react to changes in genomic integrity (Bozulic et al., 2008; Brown et al., 2015). By integrating extracellular, cytoplasmic, and nuclear cues, homeostasis can be maintained in response to a comprehensive array of physiological challenges. Until now, the biological impact of nuclear events on AKT signaling and the underlying molecular mechanisms have remained poorly understood. Here, we establish that low-level TOP2β-dependent endogenous DNA breaks regulate AKT activation. Using a viral latency model where NGF-dependent AKT activation represses virus reproduction in neurons, we demonstrate that inhibiting cellular DNA repair factors that repair TOP2β-dependent DNA breaks, such as NHEJ, TDP2, and the MRN complex, can block AKT activation and promote reactivation of latent HSV-1 infections. Unexpectedly, this process involves dynamic changes to both AKT Ser473 phosphorylation and the AKT subcellular localization; this is mediated, to a large extent, by the continued presence of extracellular NGF, DNA-PK and the AKT Ser473-specific phosphatase, PHLPP1. Thus, our study demonstrates that the DDR is tightly integrated into the growth control pathway via AKT phosphorylation dynamics, of which several nodal points can be co-opted by HSV-1 to serve as a control panel for viral latency.

### AKT dynamic localization in response to nuclear DNA damage

Surprisingly, the source of the DNA damage responsible for regulating AKT activation was not confined to exogenous agents, but instead reflected endogenous, cell intrinsic dsDNA breaks generated by TOP2βcc intermediates. Promoters of neuronal early-response genes have only been recently found to contain TOP2βcc intermediates (Madabhushi and Kim, 2018) and the idea that these might impact the AKT central signaling network is therefore unexpected. Because AKT activation occurs in proximity to the plasma membrane, phosphorylation of AKT substrates in other subcellular compartments (i.e. nucleus) was thought to rely on the subsequent diffusion of activated AKT (Calleja et al., 2007; Kunkel et al., 2005). While this conveniently accounts for the observation that AKT is activated by growth factor receptors localized at either the plasma membrane or in endosomes, its ability to target nuclear substrates would effectively uncouple AKT from its activating stimulus. Recent reports have shown that AKT activity can be confined to different activating lipid secondary messengers that could be spatially regulated throughout the cell (Ebner et al., 2017; Lučić et al., 2018). Here, we provide insight into how a cell membrane-activated AKT (NGF-stimulated) can receive an additional nuclear DNA damage signal (DNA-PK) to become fully activated and subsequently stimulate downstream cytoplasmic mTORC1 signaling (model, Figure 7). Whether AKT activation in the nucleus via phosphorylation by DNA-PK drives its nuclear function prior to its more established cytoplasmic role is presently unknown. AKT has been implicated in regulating diverse substrates that have nuclear functions, including important roles in gene expression, DDR, DNA repair, and the maintenance of genomic stability (Szymonowicz et al., 2018). For example, activated AKT in the nucleus (immediately after Ser473 phosphorylation by DNA-PK) may conceivably regulate phosphorylation of the transcription factor FOXO, which resides in the nucleus prior to AKT phosphorylation (Brunet et al., 1999; Brunet et al., 2002), or DNA repair effector proteins, such as XLF or MERIT40 (Brown et al., 2015; Liu et al., 2015). Thus, DNA damage-induced AKT activation expedites crosstalk between DNA repair/genome stability and cell growth/survival pathways (Hawkins et al., 2011), potentially impacting a diverse range of normal and pathophysiological processes, including neuronal stress responses, innate immunity, and cancer.

By directly linking perturbations in AKT phosphorylation dynamics and subcellular localization to defects in downstream mTORC1 signaling, our findings support the possibility that AKT shuttles between nuclear and cytoplasmic compartments and thereby carrying a nuclear signal (via phosphorylation) back out to the cytoplasm (inside-out signaling) to phosphorylate its downstream targets (model, Figure 7). This starkly contrasts with the typical direction of cell signaling from the cell membrane or endosomes via receptor tyrosine kinases to the nucleus to activate transcription factors, such as CREB or FOXO (Cox et al., 2008; Manning and Toker, 2017; Terenzio et al., 2017). This is reminiscent of activation of NFκB transcription factor family members resident in the cytoplasm by the nuclear DNA damage response signaling (Huang et al., 2003; McCool and Miyamoto, 2012; Wu et al., 2006). Inside-out signaling of this kind could theoretically integrate DNA damage and/ or DNA sensing with any cytoplasmic signaling event. Further investigation is required to determine the pervasiveness of this cross-talk between signaling events that spans the two compartments and the extent it regulates physiologically-relevant cell-intrinsic stress responses in different cell types and/or tissues.

By regulating AKT activation, endogenous DNA damage responses are able to control viral latency even in the presence of NGF, which before now was presumed sufficient to sustain continuous AKT and mTORC1 activation. Two of the DNA damage response factors that regulate AKT, DNA-PK and Mre11, have also been proposed to function as DNA sensors that participate in innate immune responses by sensing both microbial and non-microbial DNA (Ferguson et al., 2012; Kondo et al., 2013). This potentially provides a mechanism to integrate nuclear DNA damage responses with cytoplasmic signaling to control inflammatory responses in a variety of diseases (Crowl et al., 2017; Gao et al., 2015; Li and Chen, 2018; Sliter et al., 2018). It might further expose a strategy used by a variety of nuclear DNA viruses that subvert DNA damage response to support their long-term persistence (Anacker and Moody, 2017; Bencherit et al., 2017; Mariggiò et al., 2017; Piekna-Przybylska et al., 2017).

### Differential roles of the DDR and AKT in virus persistence and reproduction

While we establish that the DDR influences HSV-1 latency in neurons by regulating AKT, it exerts a different impact upon productive virus growth, which has been primarily investigated in nonneuronal cells. Following nuclear entry, the viral linear dsDNA genome, which contains nicks and gaps (Smith et al., 2014), triggers the host DDR (Weitzman and Fradet-Turcotte, 2018). To prevent viral genome silencing and suppress antiviral defenses, virus-encoded factors antagonize select DDR components (Lilley et al., 2011). Other facets of the DDR, like the MRN complex, localize to sub-nuclear virus replication centers, interact with viral proteins, and stimulate virus reproduction (Balasubramanian et al., 2010; Lilley et al., 2005; Lou et al., 2016; Wilkinson and Weller, 2004). In addition, whereas DNA repair by NHEJ restricts productive virus replication, the Fanconi Anemia genome stability pathway stimulates HSV-1 DNA synthesis and productive growth, in part, by antagonizing NHEJ and likely stimulating HR (Karttunen et al., 2014). Finally, instead of harnessing the DDR to activate AKT, HSV-1 expresses an AKT-like kinase during its productive growth program that shares overlapping substrate specificity with the cellular AKT (Chuluunbaatar et al., 2010; Vink et al., 2018). Thus, both AKT and the DDR plays distinct roles during virus persistence (latency) and reproduction.

### Potential new strategies for interfering with AKT activation

In our study, we showed that DSB inducers can cause transient AKT nuclear accumulation (Figures 6A and 6B). Unexpectedly, inhibition of PHLPP1 led to prolonged AKT nuclear accumulation (Figure 6C); this suggests that controlling the half-life of phosphorylated Ser473 could impact the dynamics of AKT subcellular localization and its activation kinetics. One possibility is that dephosphorylation of Ser473 controls AKT nuclear export. Alternatively, AKT could also be dephosphorylated upon its return to the cytoplasm such that an unrelated mechanism may regulate its nuclear export where it is then subsequently acted upon by PHLPP1. PHLPP1 has other substrates in addition to phosphorylated Ser473 that could perhaps regulate AKT subcellular localization and activation indirectly. Understanding the regulation and subcellular localization of PHLPP1 could shed new light into how AKT activation can be controlled by subcellular trafficking dynamics. Significantly, PHLPP1-depletion inhibited HSV-1 reactivation (Figure 5F), and as such, represents a new cellular effector that links genome integrity to AKT Ser473 phosphorylation and regulates HSV-1 latency. Antagonizing a cellular target like PHLPP1 might prevent reactivation without the concern of developing viral resistance that can undermine targeting a viral function by anti-virals (Piret and Boivin, 2016). Conversely, PHLPP1 agonists might trigger reactivation, and perhaps in combination with an anti-viral, could result in a strategy to reduce or eliminate latent viral reservoirs.

## EXPERIMENTAL PROCEDURES

### Primary Neuronal Culture

Primary superior cervical ganglion (SCG) neurons were dissected from embryonic day 21 (E21) Sprague-Dawley rat pups and cultured as described (Camarena et al., 2010; Kobayashi et al., 2012a). All protocols for isolating SCGs were approved by the Institutional Animal Care and Use Committee at NYU School of Medicine. Isolated SCGs were placed in Leibovitz’s L-15 media (Leibovitz’s L-15 with 0.4% D(+)-glucose) and incubated in collagenase (1 mg/mL, Sigma) and trypsin (0.25%, Invitrogen) at 37°C for 15 min then centrifuged for 1 min at 1000rpm. Cells were suspended in MEM medium (1x MEM, 0.4% D (+)-glucose, 2mM L-glutamine, 10% FBS) and passage through 21G and 23G needles followed by a 70-mm cell strainer then plated in 96-wells or 24-wells plates. The plates were coated with rat-tail collagen (0.66 mg/mL, Millipore) and laminin (2 μg/mL, Sigma). After 1 day culture, cells were maintained in NBM medium (neurobasal medium, 0.4% D (+)-glucose, 2mM L-glutamine, NeuroCult™ SM1 Neuronal Supplement, 50 ng/ml 2.5S NGF) supplemented with aphidicolin (5 μM, Calbiochem) and 5-fluorouracil (20 μM, Sigma) to remove proliferating cells.

### Non-neuronal Cell Culture

Rat2 (normal rat fibroblast, ATCC CRL-1764) and REF (primary embryonic fibroblast from E16 Sprague-Dawley rat pups) cells were cultured in Dulbecco’s Modified Eagle Medium (DMEM) supplemented with 10% fetal bovine serum (FBS) and 2 mM L-Glutamine.

### HSV-1 Latency Establishment and Reactivation in Primary SCG Neurons

After 6 days culture, the day before HSV-1 infection, SCG neurons were pretreated with Acyclovior (ACV) overnight. At the next day, the neuronal cultures were infected with wild-type HSV-1 (Patton strain expressing a Us11-EGFP fusion protein). Neurons were infected at an MOI of 1.5 PFU/neuron (the titer was determined in Vero cells) in NBM medium for 2 hrs. After 2 hrs, the virus was removed and the neurons were maintained in NBM medium supplemented with 50 ng/mL NGF and 100 μM ACV for at least 6 days to establish latency. After 6 days post HSV-1 infection, ACV was removed and cultures were induced to reactivate by treatment with LY294002 (20 μM, 20hrs, Calbiochem), NU7441 (1 μM, 20hrs, TOCRIS), Mirin (100 μM, 20hrs, TOCRIS), KU55933 (10 μM, Calbiochem), Etoposide (10 μM, 8hrs, Calbiochem), or Bleomycin (10 μM, 8hrs, Calbiochem). JNK inhibitor (20 μM, JNK Inhibitor II, Calbiochem) was used to block reactivation and was used for pre-treatment (6 hrs) prior to the addition of the inhibitor with other chemical inhibitors for the indicated times. Z-VAD-fmk (20 μM, 8hrs, Calbiochem) was used to block reactivation and was used concomitantly with Etoposide (8hrs) prior to detection of GFP-positive cells. Reactivation was quantified by counting the percentage of GFP-positive wells, 30 independently infected wells were analyzed for individual treatments. All the data were collected from a minimum of three separate dissection experiments (3 biological replicates). Analysis and statistical calculations were made using two-tailed unpaired Student’s t-test (Prism 7.0).

### Western Blot Analysis

Total whole cell extracts from SCGs were collected by lysis in Laemmli buffer (Bio-Rad) from approximately 50,000 neurons and fractionated on NuPAGE 4-12% Bis-Tris Protein Gels (Life Technologies). For REF and Rat2 cells, cells were lysed in lysis buffer (0.05 M Tris pH 6.8, 2% (w/v) SDS and 6% (v/v) β-mercaptoethanol) and protein concentrations were quantified using Bradford Protein Assay (Bio-Rad). Equal concentrations of protein were fractionated on NuPAGE 4-12% Bis-Tris Protein Gels. The protein was then transferred to polyvinylidene difluoride (PVDF) membranes (Millipore), blocked with 5% milk in TBST for 1 hour and incubated with primary antibody overnight at 4°C. The signals were detected by HRP-conjugated secondary antibodies and ECL reagent (PerkinElmer). Quantitative Western blotting was made using ImageJ. Primary antibodies used for Western blot analysis: anti-Ku80 (Santa Cruz Biotechnology), anti-a-tubulin (Millipore), anti-Rad50 (Santa Cruz Biotechnology), anti-Lig III (Abcam), anti-Lig IV (Novus Biologicals), anti-phospho-AKT (Ser473) (Cell Signaling Technology), anti-AKT1 (Santa Cruz Biotechnology), anti-Nbs1 (Cell Signaling Technology), anti-FLAG (Sigma-Aldrich), anti-TOP2B (Novus Biologicals), anti-phospho-Histone H2A.X (Ser139) (Cell Signaling Technology), anti-SGK1 (Abcam), anti-phospho-SGK1 (Ser422) (Abcam), anti-Rictor (Cell Signaling Technology), anti-PHLPP1 (Millipore), anti-PHLPP2 (Bethyl Laboratories), anti-TDP2 (Santa Cruz Biotechnology).

### RNA isolation and reverse-transcription quantitative PCR (RT-qPCR)

Total RNA was extracted from approximately 50,000 neurons using the Qiashedder spin column and the RNAeasy mini kit (Qiagen). Neurons were lysed in 350 μL of RLT buffer (QIAGEN) and centrifuged through a QIAGEN Shredder spin column for homogenization. Each sample were added 550 μL RNase free-water and 450 μL 100% ethanol to precipitate RNA. The samples were then applied to the RNeasy columns and washed as recommended then eluted with RNase-free water. The eluted RNA was treated with DNase I (New England Biolabs) for 10 min at 37°C and the DNase I was heat inactivated at 75°C for 10 min in the presence of 2 mM EDTA. cDNA was generated using qScript™ cDNA SuperMix (QuantaBio). Quantitative realtime PCR (qRT-PCR) analysis was performed using SYBR™ Green PCR Master Mix (Applied Biosystem) and a StepOnePlus Real-Time PCR System (Applied Biosystem). Relative expression levels of viral mRNAs were normalized to 18S rRNA (Camarena et al., 2010; Kobayashi et al., 2012b), statistical calculations were made using two-tailed paired students T-test (Prism 7.0).

### Immunofluorescence

Neurons were seeded at a density of 2×10^4^ cells/well onto coverslips pre-coated with poly-D lysine and laminin. After treatments, cultures were washed twice with PBS and fixed with 4% paraformaldehyde in PBS for 15 min at RT then permeabilized with 0.25% Triton X-100 in PBS. To detect AKT1 and γ-H2Ax nuclear localization, cells were pre-permeabilized with Scully buffer (0.5% Triton X-100, 20 mM HEPES-KOH, pH 7.4, 50 mM NaCl, 3 mM MgCl2, 300 mM sucrose) for 1 min and fixed with 4% paraformaldehyde in PBS for 15 min at RT. After blocking in 2% BSA in PBS for 20 min, cells were incubated with primary antibodies for 3 hours at RT and detected using anti-mouse Alexa Fluor^®^ 488, anti-rabbit Alexa Fluor^®^ 488 or anti-rabbit Alexa Fluor^®^ 546 for 2 hours at RT. Nuclei were stained with Hoechst 33258 (Life Technologies). Images were acquired by Zeiss LSM 700 confocal microscope software and were processed using ImageJ. Primary antibodies used for immunofluorescence studies: anti-GFP (Santa Cruz Biotechnology), anti-phospho-Histone H2A.X (Ser139) (Cell Signaling Technology), anti-β3-Tubulin (Cell Signaling Technology), anti-AKT1 (Santa Cruz Biotechnology), anti-mTOR (Cell Signaling Technology), anti-TSC2 (Cell Signaling Technology).

### Preparation of HSV-1 stocks and plaque assay

HSV-1 stocks were amplified and titered on Vero cell monolayers by standard methods. Vero cells were infected (MOI=0.01) and maintained in Dulbecco’s modified Eagle’s medium supplemented with 10% FBS and 2 mM L-glutamine at 37°C for 2–3 days then harvest by freeze-thaw lysis and sonication. Infectious titers were determined by plaque assay using Vero cells. For plaque assays, uninfected Vero cells were seeded in 6-well plates and incubated with serial diluted virus for 2 h at 37°C then overlaid with 0.5% agarose in MEM supplemented with 1% FBS. After incubation for 2-3 days at 37°C, cells were fixed with 10% TCA for 10 min and stained with 1% crystal violet to visualize the plaques.

### Preparation of lentiviral stocks and lentivirus infection of SCG neurons

Lentiviruses expressing shRNAs (Sigma-Aldrich Mission mouse or human Lentiviral shRNA libraries) or Flag-Rheb (Q64L) were generated using the 293LTV packaging cell line. The envelop plasmid (pMD2.G), packaging plasmid (psPAX2), and pLKO.1 or pcw107 control vectors were cotransfected into 293LTV cells, and viral particles were harvested 48 hr posttransfection. Lentiviral particles (MOI=10) were added to neurons overnight. For expression of the Rheb (Q64L) mutant, latently-infected neurons were transduced with Rheb (Q64L) or control pcw107 lentiviral particles overnight. After two days, ACV was removed and cultures were induced to reactivate by treatment with LY294002 (20 μM), NU7441 (1 μM), or Mirin (100 μM). 20 hours later, mRNA was prepared and analyzed by qRT-PCR. Flag-Rheb (Q64L)-pcw107 was a gift from David Sabatini and Kris Wood (Addgene plasmid #64607).

### ICE (In vivo Complex of Enzyme) assay

TOP2β-DNA complexes were isolated from approximately 600,000 SCG neurons using the ICE (In Vivo Complex of Enzyme) assay kit (TopoGEN). All procedures were performed as instructions. Cells were washed in PBS twice and lysed in lysis buffer. Genomic DNA was precipitated by 100% ethanol and then centrifuged. Pellets were washed with 75% ethanol and resuspended in buffer D (provided from the kit). DNA concentration was determined by nanodrop (Thermo Scientific) then applied to a methanol-pretreated PVDF membrane using a Bio-Dot apparatus (Bio-Rad). The membranes were then probed for TOP2β using standard western blotting procedures.

### Statistical analysis

All statistical analysis was performed using Prism software (7.0). Number of biological repeats of experiments and statistical significance are indicated in the corresponding figure legends. P values equal to or less than 0.05 were considered significant, asterisks denote statistical significance (*, p < 0.05; **, p < 0.01; ***, p < 0.001; ****, p < 0.0001). P-values are calculated using two-tailed unpaired Student’s t-test. P values greater than 0.05 were not significant (NS).

## KEY RESOURCES TABLE

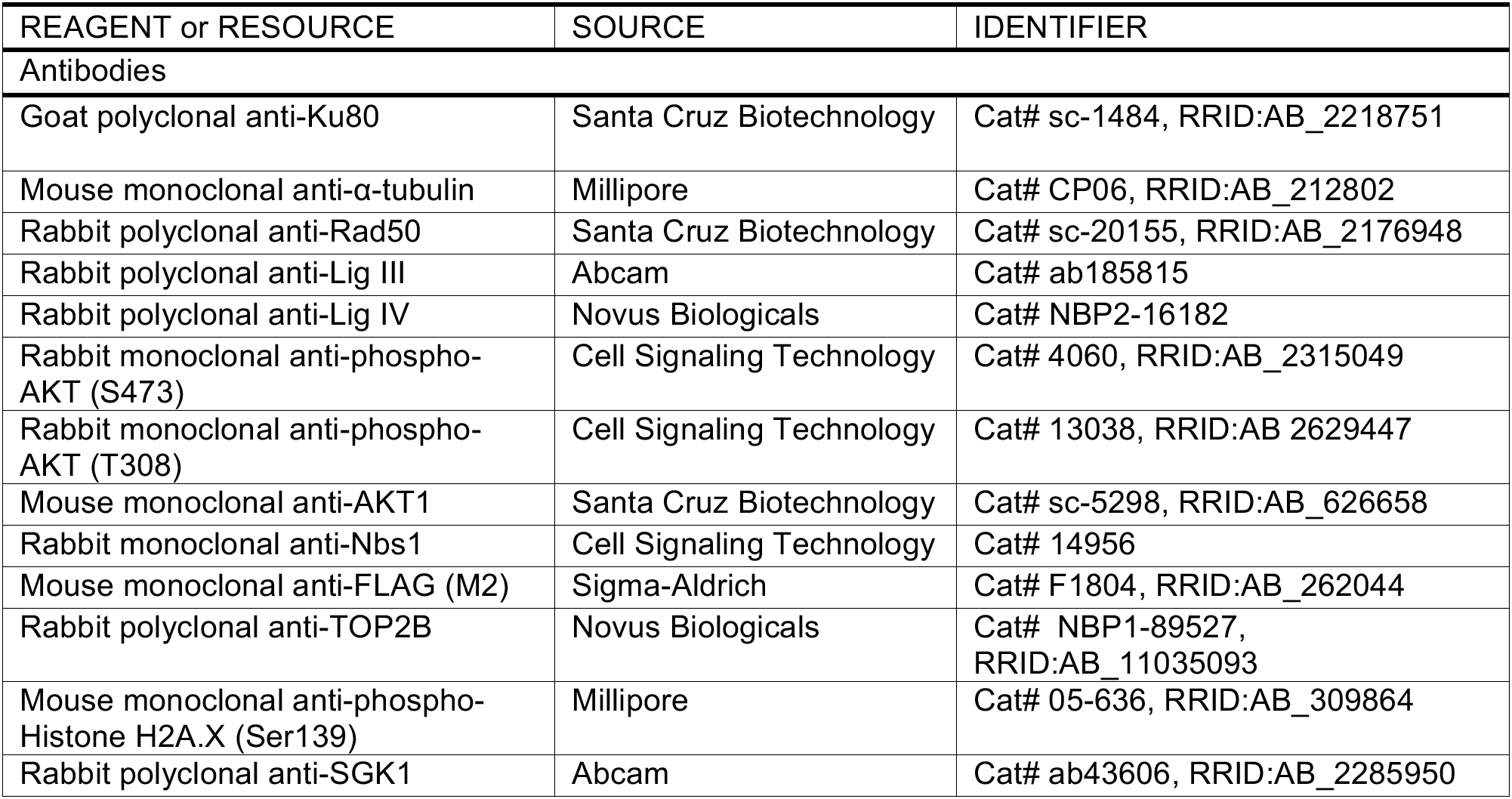
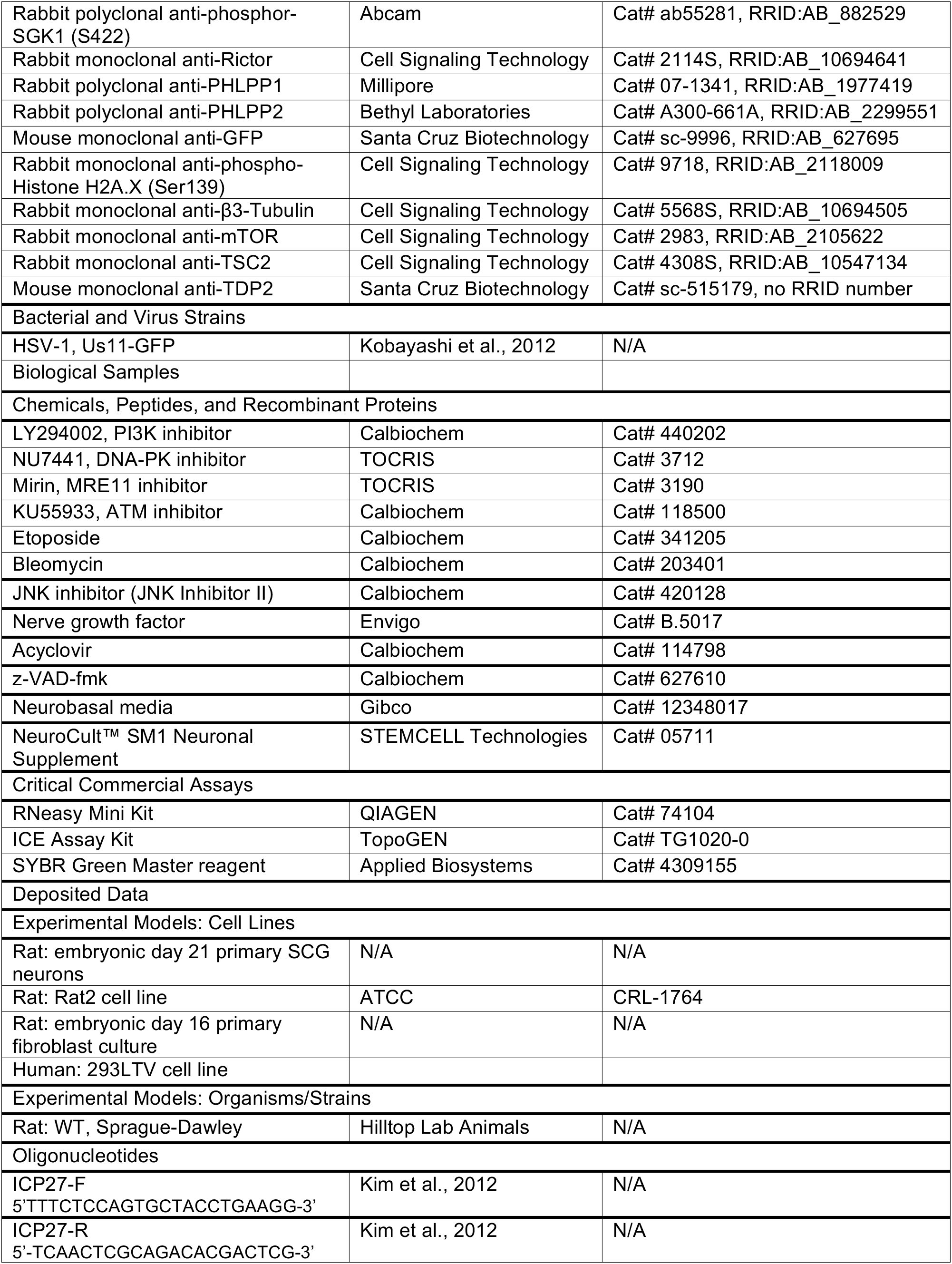
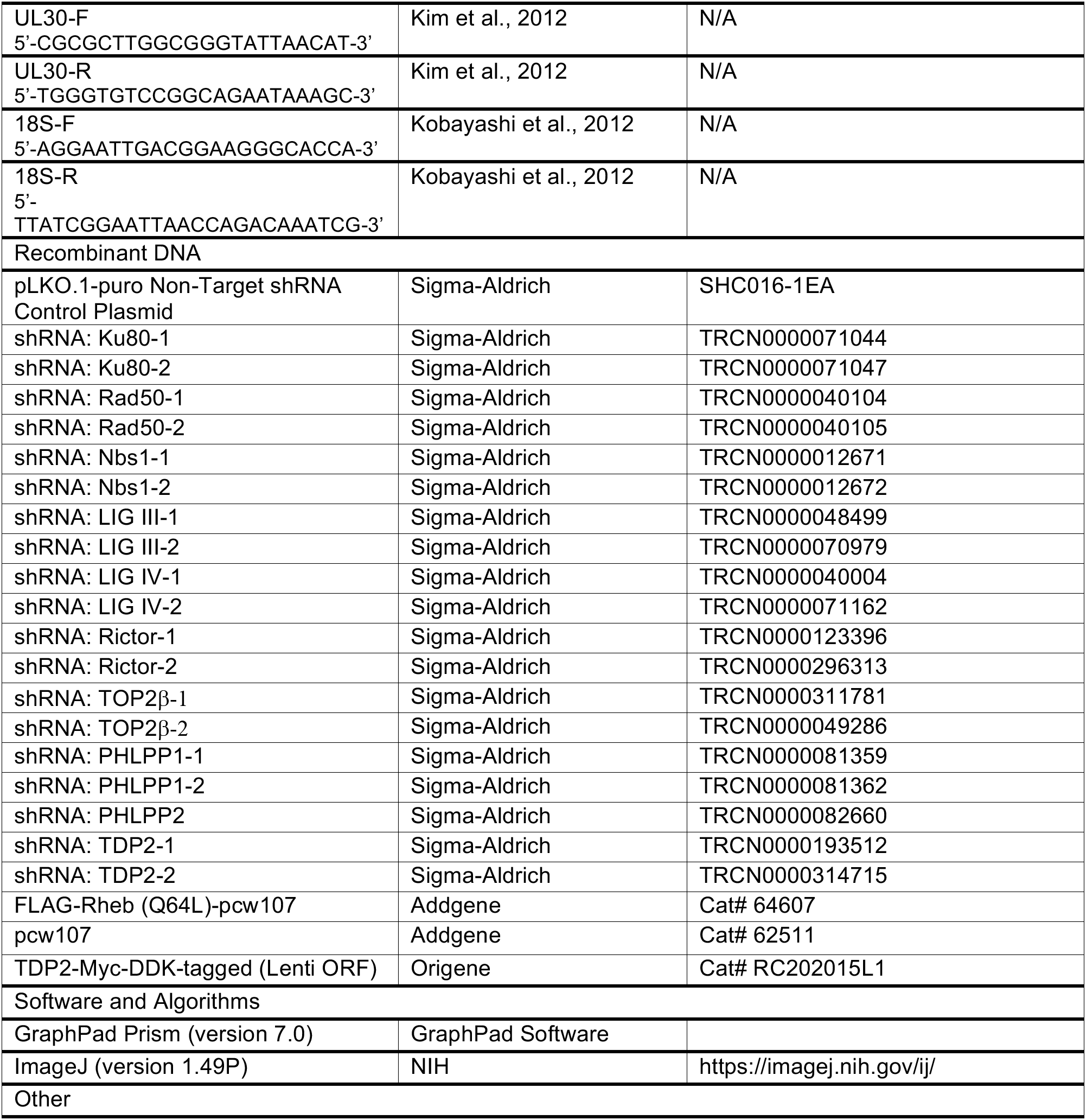

## AUTHOR CONTRIBUTIONS

H.H., I.M., and T.T.H. designed the study, H.H. performed the experiments with technical support from L.S. and M.K., M.C. provided critical reagents and equipment for the study, H.H., A.W., I.M. and T.T.H analyzed the data, and H.H., M.C., A.W., I.M. and T.T.H. wrote the manuscript.

## ACKNOWLEDGEMENTS

We thank J. Linderman for advice and guidance on primary neuronal culture and preparation of HSV-1 virus stocks, and members of the Huang, Mohr, Wilson and Chao labs for critical discussions. This work was supported by NIH grants GM107257 (T.T.H.), GM056927, AI073898 (I.M.), AI130618 (A.W.), and R56 NS021072 (M.V.C.), and funds from the V Foundation for BRCA Cancer Research to T.T.H.

---

**Supplementary Figure 1.**
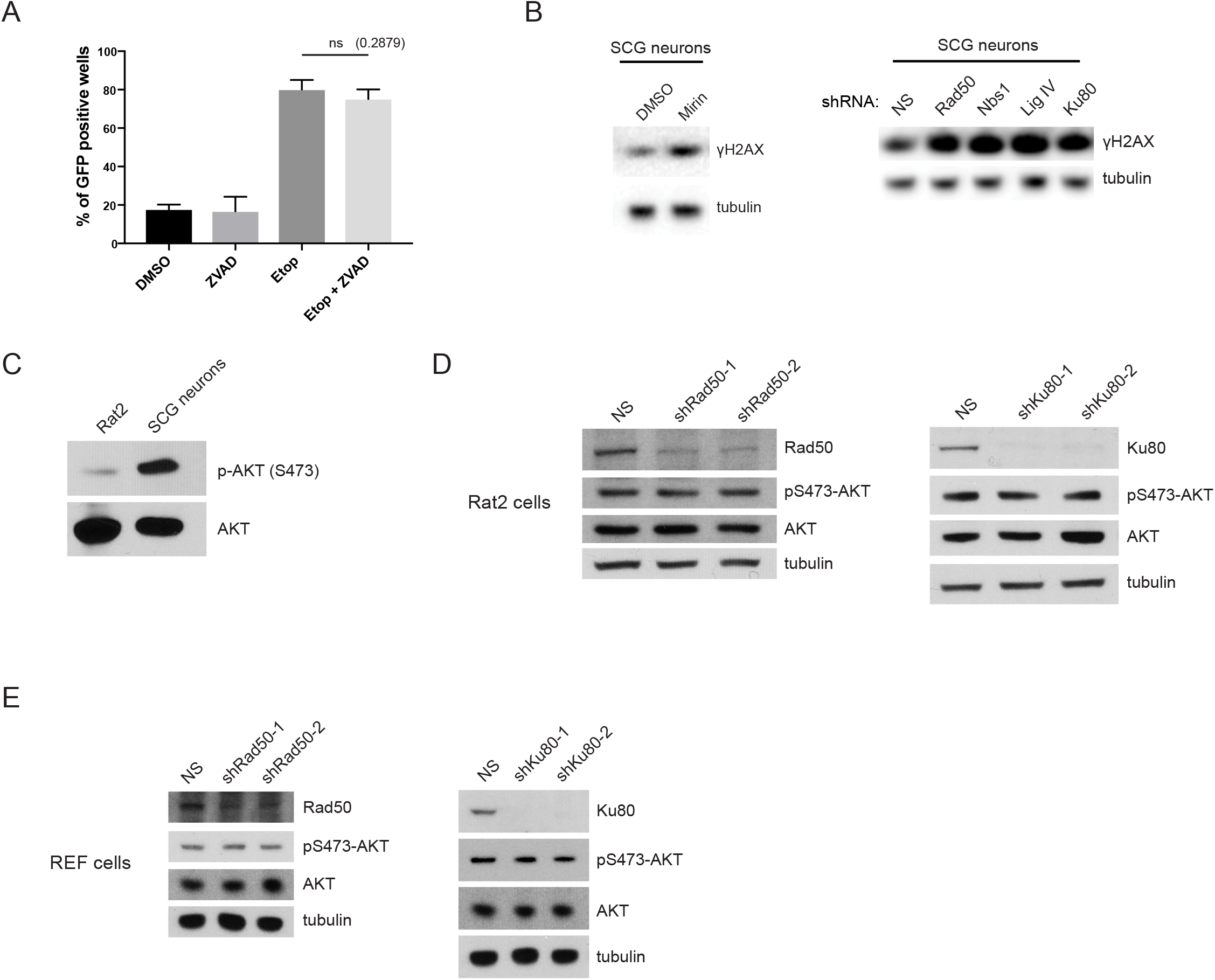
The loss of MRN or NHEJ factors do not affect Akt phosphorylation in non-neuronal rat cells. (A) Neuronal apoptosis is not required for HSV-1 reactivation after Etoposide treatment. HSV-1 latently-infected SCG neurons were treated with Etoposide (10μM) or DMSO in the presence or absence of 20 μM Z-VAD-fmk. Reactivation assay showing the percentages of wells containing GFP-positive cells deteced by fluorescent microscopy of living neurons at day 3 post-treatment. (B) Inhibiting Mre11 or knockdown of different NHEJ or MRN complex components elevates γH2AX levels in SCG neurons. Western blot analysis of uninfected SCG neurons treated with Mirin (100 μM) or DMSO control for 8 hrs (left panel), or untreated SCG neurons lentivirally-transduced with different shRNAs for 5 days (right panel). (C) Western blot analysis comparing Akt activation in Rat2 normal fibroblasts vs SCG neurons. (D-E) Lentiviral-expressed shRNAs against Rad50 or Ku80 in either the Rat2 fibroblast cell line (D) or primary REFs (E) and analyzed by Western blot with the indicated antibodies.

**Supplementary Figure 2.**
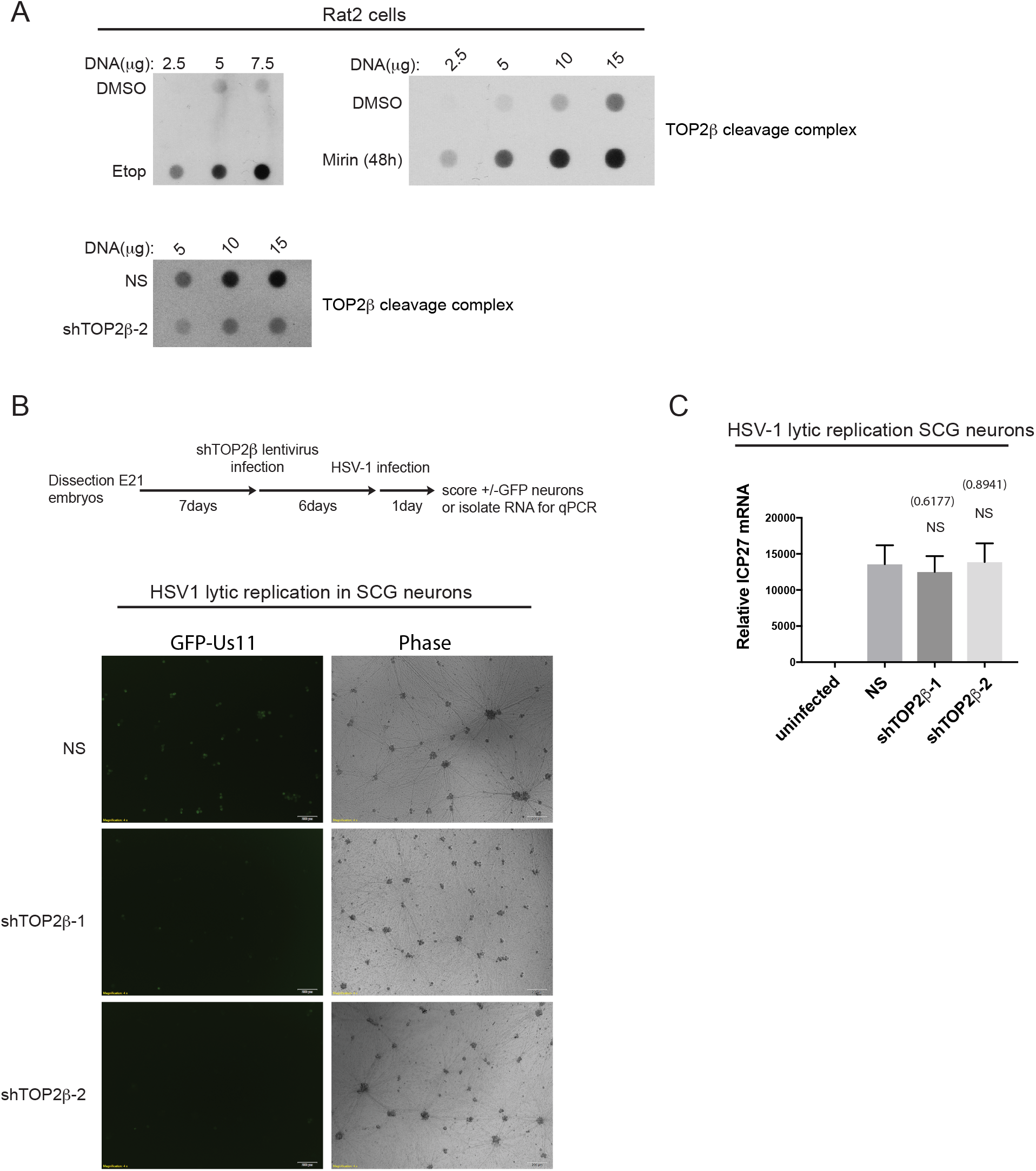
Detection of TOP2βcc upon Etoposide or Mirin treatments in Rat2 cells. (A) Dot blot analysis of TOP2βcc levels in Rat2 cells treated with either DMSO control or Etoposide (10μM, 20 hrs) or Mirin (100μM, 48hrs), or with an shRNA against TOP2β. Different amounts of genomic DNA were prepared as indicated. (B) Schematic for HSV-1 lytic infection detection of GFP-positive SCG neurons (no acyclovir treatment). Live-cell imaging of GFP+ fluorescence in SCG neurons undergoing lytic replication. SCG neurons were depleted for TOP2β for 6 days prior to HSV-1 infection (1 day). Loss of TOP2β compromised HSV-1 lytic replication due to the absence of GFP-Us11 accumulation. (C) Induction of ICP27 viral mRNA (qRT-PCR) was unaffected by the loss of TOP2β in SCG neurons under lytic infection (3 biological replicates with -/+ s.e.m., NS is not significant, p > 0.05).

**Supplementary Figure 3.**
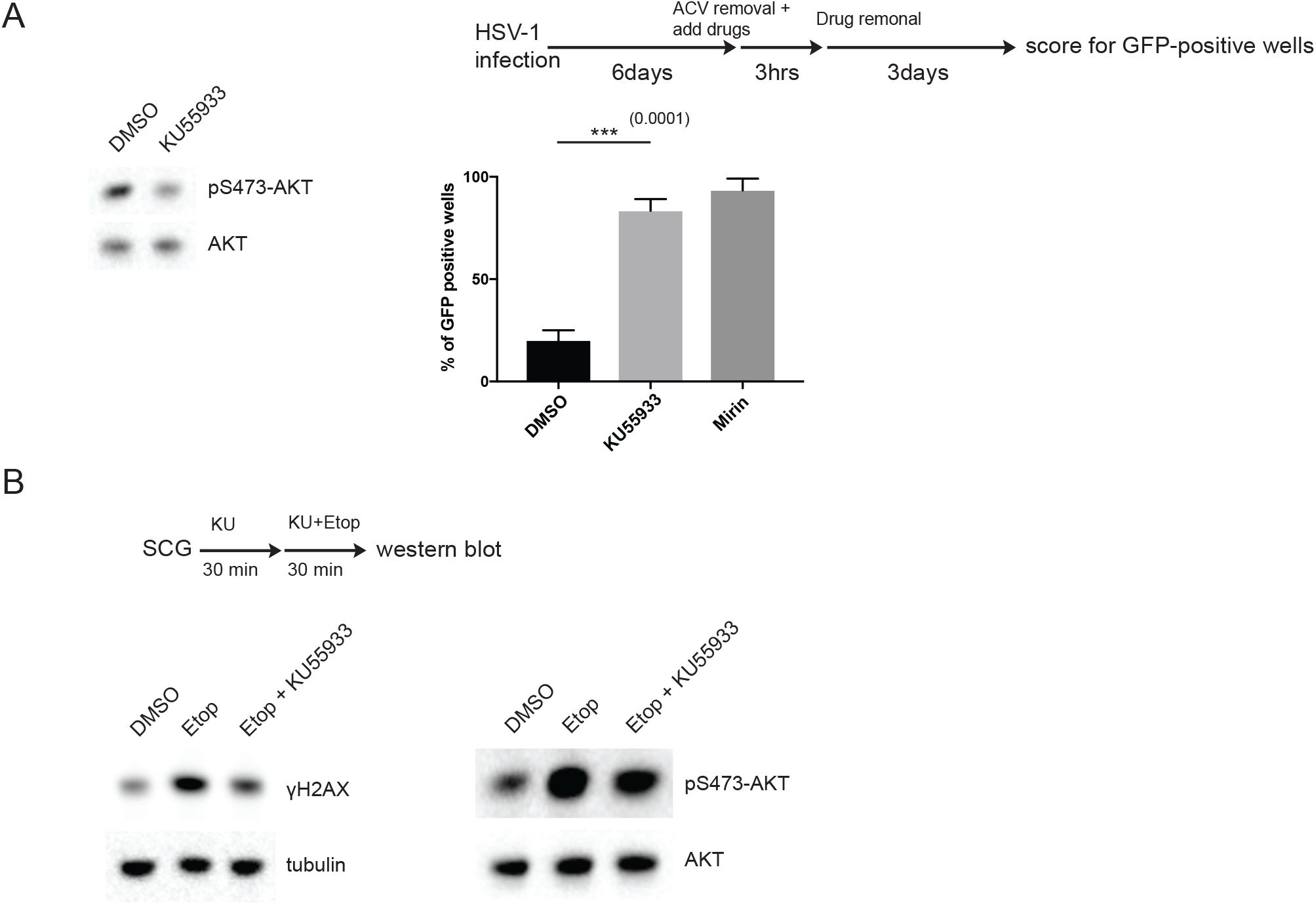
Treatment of SCG neurons with ATM inhibitor causes HSV-1 reactivation. (A) Uninfected SCG neurons treated with either DMSO or KU55933 (ATM inhibitor, 10μM for 3 hrs) and analyzed by Western blot for its effect on Akt Ser473 phosphorylation (left panel). HSV-1 latently-infected SCG neurons were treated with either KU55933 or Mirin as indicated in the schematics (right panel). The number of GFP+ wells from a 96-well plate were scored for with +/− s.e.m. for 3 biological replicates and p-values calculated using two-tailed unpaired Student’s t-test. (B) Modest effect on H2AX and Akt phosphorylation after KU55933 treatment in response to Etoposide (see treatment schematics). Western blot analysis was performed to probe the indicated proteins.

**Supplementary Figure 4.**
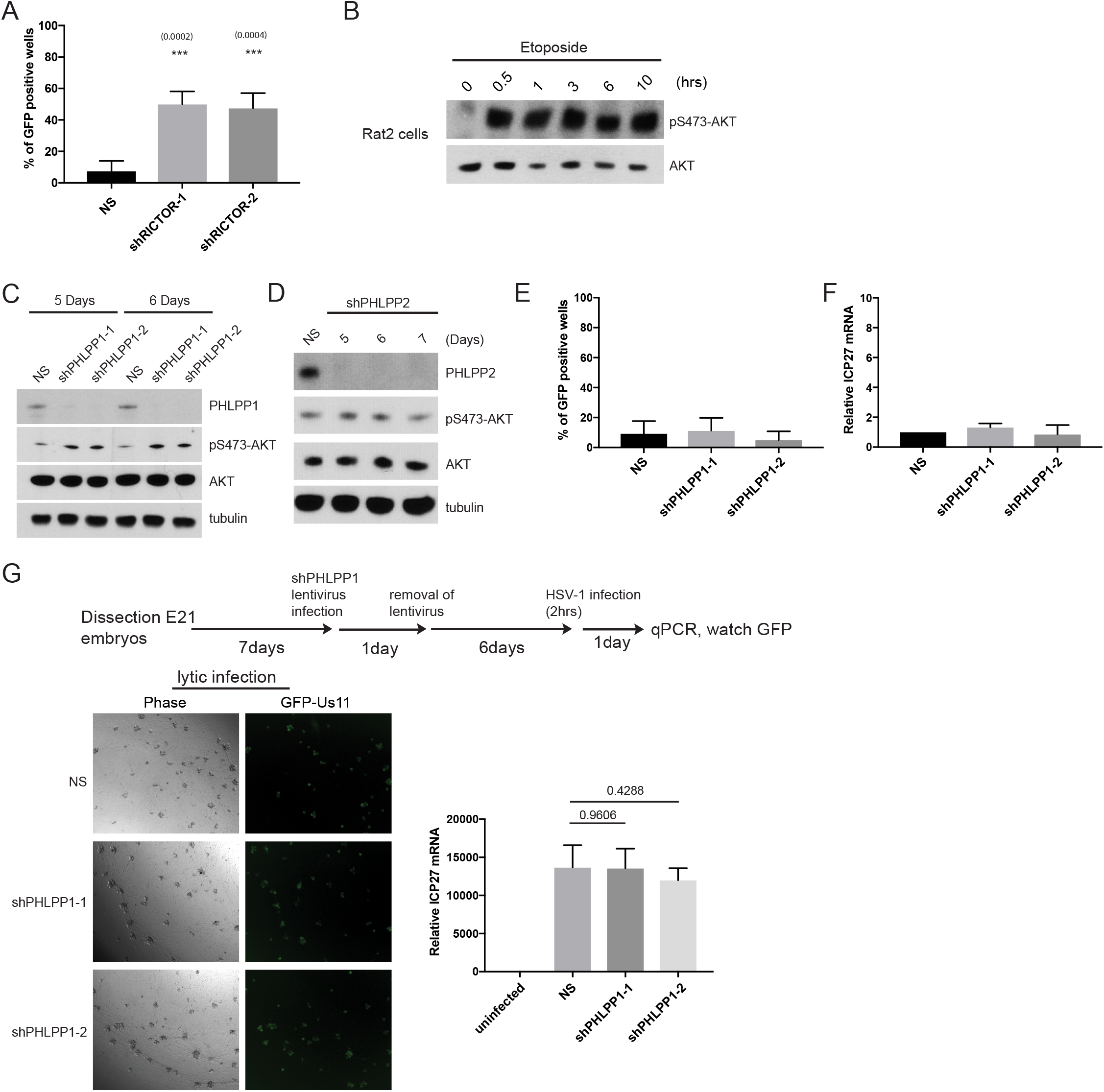
Loss of RICTOR, but not PHLPP1, caused an increase in spontaneous HSV-1 reactivation. (A) Depletion of RICTOR using lentiviral-delivered shRNAs in latently-infected SCG neurons. Number of GFP+ wells from a 96-well plate were scored for with +/− s.e.m. for 4 biological replicates and p-values calculated using two-tailed unpaired Student’s t-test. (B) Time-course study with Etoposide treatment in Rat2 cells for Akt phosphorylation by Western blot analysis. (C-D) Depletion of either PHLPP1 using two different shRNAs or PHLPP2 in SCG neurons were analyzed after different days post-lentiviral infection. Effects on Akt phosphorylation were displayed by Western blot analysis using the indicated antibodies. (E-F) Depletion of PHLPP1 using lentiviral-delivered shRNAs (6 days) in latently-infected SCG neurons were scored for spontaneous HSV-1 reactivation by GFP+ wells (E), or with qRT-PCR for elevated levels of ICP27 viral mRNA (F) after 1 day. (G) Depletion of PHLPP1 did not affect HSV-1 lytic infection in SCG neurons. SCG neurons were depleted of PHLPP1 by shRNAs and then infected with HSV-1 in the absence of acyclovir (1 day) to induce lytic infection (see schematics). Levels of GFP-positive neurons (left panel) and ICP27 viral mRNA induction (right panel) observed were comparable between NS and PHLPP1 shRNA-treated neurons (not significant, P value greater than 0.05).

**Supplementary Figure 5.**
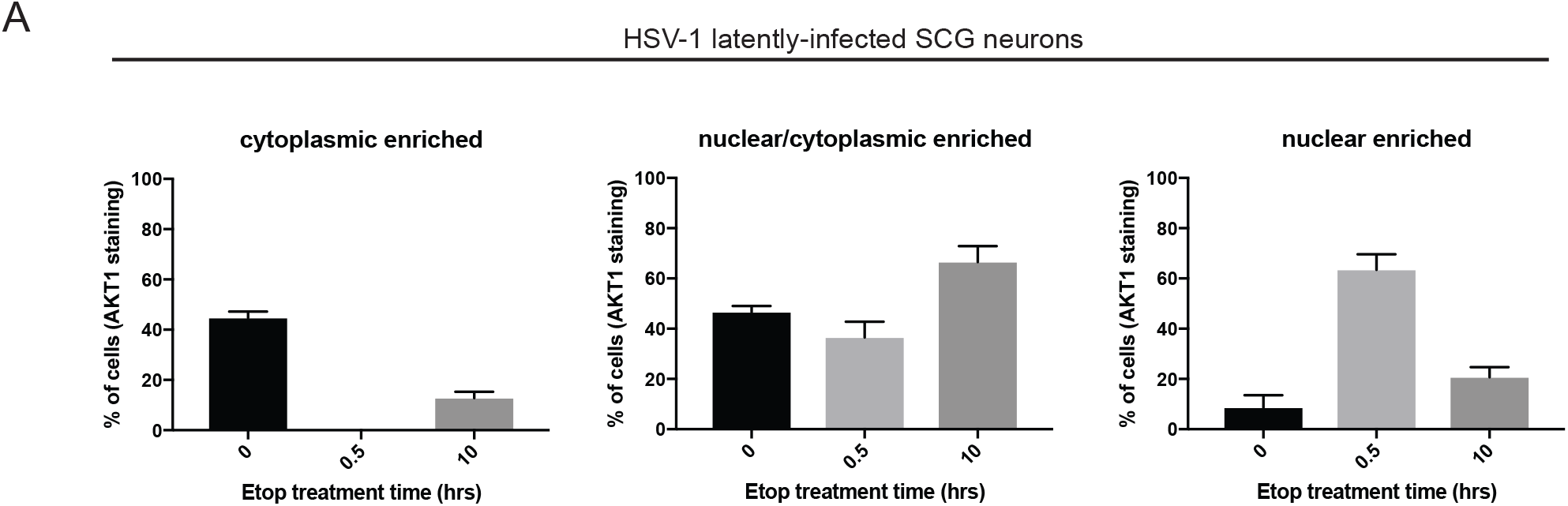
Akt subcellular localization in response to Etoposide treatment are comparable in both HSV-1 latently-infected and uninfected SCG neurons. (A) Cultured SCG neurons that are latently infected with HSV-1 are immunostained with Akt1 antibody were left either untreated (0h) or treated with Etoposide for indicated times and imaged by confocal microscopy. Nuclear DNA was visualized using Hoechst stain to determine the boundaries of the nucleus/cytoplasm. The percentage of cells with either cytoplasmic, nuclear/cytoplasmic, or nuclear-enriched Akt1 subcellular localization were quantified and displayed as bar graphs with mean +/− s.e.m. for 3 biological replicates. These results are similar to the uninfected SCG neurons as shown in main Figure 5A.

## REFERENCES

Anacker, D.C., and Moody, C.A. (2017). Modulation of the DNA damage response during the life cycle of human papillomaviruses. Virus Res 231, 41–49.

Aparicio, T., Baer, R., Gottesman, M., and Gautier, J. (2016). MRN, CtIP, and BRCA1 mediate repair of topoisomerase II-DNA adducts. J Cell Biol 212, 399–408.

Balasubramanian, N., Bai, P., Buchek, G., Korza, G., and Weller, S.K. (2010). Physical interaction between the herpes simplex virus type 1 exonuclease, UL12, and the DNA double-strand break-sensing MRN complex. J Virol 84, 12504–12514.

Benboudjema, L., Mulvey, M., Gao, Y., Pimplikar, S.W., and Mohr, I. (2003). Association of the herpes simplex virus type 1 Us11 gene product with the cellular kinesin light-chain-related protein PAT1 results in the redistribution of both polypeptides. J Virol 77, 9192–9203.

Bencherit, D., Remy, S., Le Vern, Y., Vychodil, T., Bertzbach, L.D., Kaufer, B.B., Denesvre, C., and Trapp-Fragnet, L. (2017). Induction of DNA Damages upon Marek’s Disease Virus Infection: Implication in Viral Replication and Pathogenesis. J Virol 91.

Bozulic, L., Surucu, B., Hynx, D., and Hemmings, B.A. (2008). PKBalpha/Akt1 acts downstream of DNA-PK in the DNA double-strand break response and promotes survival. Mol Cell 30, 203–213.

Brognard, J., Sierecki, E., Gao, T., and Newton, A.C. (2007). PHLPP and a second isoform,PHLPP2, differentially attenuate the amplitude of Akt signaling by regulating distinct Akt isoforms. Mol Cell 25, 917–931.

Brown, K.K., Montaser-Kouhsari, L., Beck, A.H., and Toker, A. (2015). MERIT40 Is an Akt Substrate that Promotes Resolution of DNA Damage Induced by Chemotherapy. Cell Rep 11, 1358–1366.

Brunet, A., Bonni, A., Zigmond, M.J., Lin, M.Z., Juo, P., Hu, L.S., Anderson, M.J., Arden, K.C., Blenis, J., and Greenberg, M.E. (1999). Akt promotes cell survival by phosphorylating and inhibiting a Forkhead transcription factor. Cell 96, 857–868.

Brunet, A., Kanai, F., Stehn, J., Xu, J., Sarbassova, D., Frangioni, J.V., Dalal, S.N., DeCaprio, J.A., Greenberg, M.E., and Yaffe, M.B. (2002). 14-3-3 transits to the nucleus and participates in dynamic nucleocytoplasmic transport. J Cell Biol 156, 817–828.

Calleja, V., Alcor, D., Laguerre, M., Park, J., Vojnovic, B., Hemmings, B.A., Downward, J., Parker, P.J., and Larijani, B. (2007). Intramolecular and intermolecular interactions of protein kinase B define its activation in vivo. PLoS Biol 5, e95.

Camarena, V., Kobayashi, M., Kim, J.Y., Roehm, P., Perez, R., Gardner, J., Wilson, A.C., Mohr, I., and Chao, M.V. (2010). Nature and duration of growth factor signaling through receptor tyrosine kinases regulates HSV-1 latency in neurons. Cell Host Microbe 8, 320–330.

Chang, H.H.Y., Pannunzio, N.R., Adachi, N., and Lieber, M.R. (2017). Non-homologous DNA end joining and alternative pathways to double-strand break repair. Nat Rev Mol Cell Biol 18, 495–506.

Chuluunbaatar, U., Roller, R., Feldman, M.E., Brown, S., Shokat, K.M., and Mohr, I. (2010). Constitutive mTORC1 activation by a herpesvirus Akt surrogate stimulates mRNA translation and viral replication. Genes Dev 24, 2627–2639.

Cliffe, A.R., Arbuckle, J.H., Vogel, J.L., Geden, M.J., Rothbart, S.B., Cusack, C.L., Strahl, B.D., Kristie, T.M., and Deshmukh, M. (2015). Neuronal Stress Pathway Mediating a Histone Methyl/Phospho Switch Is Required for Herpes Simplex Virus Reactivation. Cell Host Microbe 18, 649–658.

Cortes Ledesma, F., El Khamisy, S.F., Zuma, M.C., Osborn, K., and Caldecott, K.W. (2009). A human 5’-tyrosyl DNA phosphodiesterase that repairs topoisomerase-mediated DNA damage. Nature 461, 674–678.

Cox, L.J., Hengst, U., Gurskaya, N.G., Lukyanov, K.A., and Jaffrey, S.R. (2008). Intra-axonal translation and retrograde trafficking of CREB promotes neuronal survival. Nat Cell Biol 10, 149–159.

Crowl, J.T., Gray, E.E., Pestal, K., Volkman, H.E., and Stetson, D.B. (2017). Intracellular Nucleic Acid Detection in Autoimmunity. Annu Rev Immunol 35, 313–336.

Du, T., Han, Z., Zhou, G., and Roizman, B. (2015). Patterns of accumulation of miRNAs encoded by herpes simplex virus during productive infection, latency, and on reactivation. Proc Natl Acad Sci U S A 112, E49–55.

Ebner, M., Lučić, I., Leonard, T.A., and Yudushkin, I. (2017). PI(3,4,5)P. Mol Cell 65, 416–431.e416.

Ferguson, B.J., Mansur, D.S., Peters, N.E., Ren, H., and Smith, G.L. (2012). DNA-PK is a DNA sensor for IRF-3-dependent innate immunity. Elife 1, e00047.

Fraser, M., Harding, S.M., Zhao, H., Coackley, C., Durocher, D., and Bristow, R.G. (2011). MRE11 promotes AKT phosphorylation in direct response to DNA double-strand breaks. Cell Cycle 10, 2218–2232.

Gao, D., Li, T., Li, X.D., Chen, X., Li, Q.Z., Wight-Carter, M., and Chen, Z.J. (2015). Activation of cyclic GMP-AMP synthase by self-DNA causes autoimmune diseases. Proc Natl Acad Sci U S A 112, E5699–5705.

Gao, R., Schellenberg, M.J., Huang, S.Y., Abdelmalak, M., Marchand, C., Nitiss, K.C., Nitiss, J.L., Williams, R.S., and Pommier, Y. (2014). Proteolytic degradation of topoisomerase II (Top2) enables the processing of Top2·DNA and Top2·RNA covalent complexes by tyrosyl-DNA-phosphodiesterase 2 (TDP2). J Biol Chem 289, 17960–17969.

Gao, T., Furnari, F., and Newton, A.C. (2005). PHLPP: a phosphatase that directly dephosphorylates Akt, promotes apoptosis, and suppresses tumor growth. Mol Cell 18, 13–24.

Glaser, R., and Kiecolt-Glaser, J.K. (2005). Stress-induced immune dysfunction: implications for health. Nat Rev Immunol 5, 243–251.

Glebova, N.O., and Ginty, D.D. (2005). Growth and survival signals controlling sympathetic nervous system development. Annu Rev Neurosci 28, 191–222.

Hawkins, A.J., Golding, S.E., Khalil, A., and Valerie, K. (2011). DNA double-strand break - induced pro-survival signaling. Radiother Oncol 101, 13–17.

Hoa, N.N., Shimizu, T., Zhou, Z.W., Wang, Z.Q., Deshpande, R.A., Paull, T.T., Akter, S., Tsuda, M., Furuta, R., Tsutsui, K., et al. (2016). Mre11 Is Essential for the Removal of Lethal Topoisomerase 2 Covalent Cleavage Complexes. Mol Cell 64, 580–592.

Hoeijmakers, J.H. (2001). Genome maintenance mechanisms for preventing cancer. Nature 411, 366–374.

Huang, T.T., Wuerzberger-Davis, S.M., Wu, Z.H., and Miyamoto, S. (2003). Sequential modification of NEMO/IKKgamma by SUMO-1 and ubiquitin mediates NF-kappaB activation by genotoxic stress. Cell 115, 565–576.

Karttunen, H., Savas, J.N., McKinney, C., Chen, Y.H., Yates, J.R., Hukkanen, V., Huang, T.T., and Mohr, I. (2014). Co-opting the Fanconi Anemia Genomic Stability Pathway Enables Herpesvirus DNA Synthesis and Productive Growth. Mol Cell.

Kim, J.Y., Mandarino, A., Chao, M.V., Mohr, I., and Wilson, A.C. (2012). Transient reversal of episome silencing precedes VP16-dependent transcription during reactivation of latent HSV-1 in neurons. PLoS Pathog 8, e1002540.

Knipe, D.M., and Cliffe, A. (2008). Chromatin control of herpes simplex virus lytic and latent infection. Nat Rev Microbiol 6, 211–221.

Kobayashi, M., Kim, J.Y., Camarena, V., Roehm, P.C., Chao, M.V., Wilson, A.C., and Mohr, I. (2012a). A primary neuron culture system for the study of herpes simplex virus latency and reactivation. J Vis Exp.

Kobayashi, M., Wilson, A.C., Chao, M.V., and Mohr, I. (2012b). Control of viral latency in neurons by axonal mTOR signaling and the 4E-BP translation repressor. Genes Dev 26, 1527–1532.

Kondo, T., Kobayashi, J., Saitoh, T., Maruyama, K., Ishii, K.J., Barber, G.N., Komatsu, K., Akira, S., and Kawai, T. (2013). DNA damage sensor MRE11 recognizes cytosolic double-stranded DNA and induces type I interferon by regulating STING trafficking. Proc Natl Acad Sci U S A 110, 2969–2974.

Kunkel, M.T., Ni, Q., Tsien, R.Y., Zhang, J., and Newton, A.C. (2005). Spatio-temporal dynamics of protein kinase B/Akt signaling revealed by a genetically encoded fluorescent reporter. J Biol Chem 280, 5581–5587.

Li, T., and Chen, Z.J. (2018). The cGAS-cGAMP-STING pathway connects DNA damage to inflammation, senescence, and cancer. J Exp Med 215, 1287–1299.

Lilley, C.E., Carson, C.T., Muotri, A.R., Gage, F.H., and Weitzman, M.D. (2005). DNA repair proteins affect the lifecycle of herpes simplex virus 1. Proc Natl Acad Sci U S A 102, 5844–5849.

Lilley, C.E., Chaurushiya, M.S., Boutell, C., Everett, R.D., and Weitzman, M.D. (2011). The intrinsic antiviral defense to incoming HSV-1 genomes includes specific DNA repair proteins and is counteracted by the viral protein ICP0. PLoS Pathog 7, e1002084.

Linderman, J.A., Kobayashi, M., Rayannavar, V., Fak, J.J., Darnell, R.B., Chao, M.V., Wilson, A.C., and Mohr, I. (2017). Immune Escape via a Transient Gene Expression Program Enables Productive Replication of a Latent Pathogen. Cell Rep 18, 1312–1323.

Liu, P., Gan, W., Guo, C., Xie, A., Gao, D., Guo, J., Zhang, J., Willis, N., Su, A., Asara, J.M., et al. (2015). Akt-mediated phosphorylation of XLF impairs non-homologous end-joining DNA repair. Mol Cell 57, 648–661.

Looker, K.J., Magaret, A.S., May, M.T., Turner, K.M., Vickerman, P., Gottlieb, S.L., and Newman, L.M. (2015). Global and Regional Estimates of Prevalent and Incident Herpes Simplex Virus Type 1 Infections in 2012. PLoS One 10, e0140765.

Lou, D.I., Kim, E.T., Meyerson, N.R., Pancholi, N.J., Mohni, K.N., Enard, D., Petrov, D.A., Weller, S.K., Weitzman, M.D., and Sawyer, S.L. (2016). An Intrinsically Disordered Region of the DNA Repair Protein Nbs1 Is a Species-Specific Barrier to Herpes Simplex Virus 1 in Primates. Cell Host Microbe 20, 178–188.

Lučić, I., Rathinaswamy, M.K., Truebestein, L., Hamelin, D.J., Burke, J.E., and Leonard, T.A. (2018). Conformational sampling of membranes by Akt controls its activation and inactivation. Proc Natl Acad Sci U S A 115, E3940–E3949.

Madabhushi, R., Gao, F., Pfenning, A.R., Pan, L., Yamakawa, S., Seo, J., Rueda, R., Phan, T.X., Yamakawa, H., Pao, P.C., et al. (2015). Activity-Induced DNA Breaks Govern the Expression of Neuronal Early-Response Genes. Cell 161, 1592–1605.

Madabhushi, R., and Kim, T.K. (2018). Emerging themes in neuronal activity-dependent gene expression. Mol Cell Neurosci 87, 27–34.

Madabhushi, R., Pan, L., and Tsai, L.H. (2014). DNA damage and its links to neurodegeneration. Neuron 83, 266–282.

Manning, B.D., and Toker, A. (2017). AKT/PKB Signaling: Navigating the Network. Cell 169, 381–405.

Mariggiò, G., Koch, S., Zhang, G., Weidner-Glunde, M., Rückert, J., Kati, S., Santag, S., and Schulz, T.F. (2017). Kaposi Sarcoma Herpesvirus (KSHV) Latency-Associated Nuclear Antigen (LANA) recruits components of the MRN (Mre11-Rad50-NBS1) repair complex to modulate an innate immune signaling pathway and viral latency. PLoS Pathog 13, e1006335.

Martz, C.A., Ottina, K.A., Singleton, K.R., Jasper, J.S., Wardell, S.E., Peraza-Penton, A., Anderson, G.R., Winter, P.S., Wang, T., Alley, H.M., et al. (2014). Systematic identification of signaling pathways with potential to confer anticancer drug resistance. Sci Signal 7, ra121.

Mateos-Gomez, P.A., Gong, F., Nair, N., Miller, K.M., Lazzerini-Denchi, E., and Sfeir, A. (2015). Mammalian polymerase θ promotes alternative NHEJ and suppresses recombination. Nature 518, 254–257.

McCool, K.W., and Miyamoto, S. (2012). DNA damage-dependent NF-κB activation: NEMO turns nuclear signaling inside out. Immunol Rev 246, 311–326.

McKinnon, P.J. (2013). Maintaining genome stability in the nervous system. Nat Neurosci 16, 1523–1529.

Nitiss, J.L. (2009a). DNA topoisomerase II and its growing repertoire of biological functions. Nat Rev Cancer 9, 327–337.

Nitiss, J.L. (2009b). Targeting DNA topoisomerase II in cancer chemotherapy. Nat Rev Cancer 9, 338–350.

Piekna-Przybylska, D., Sharma, G., Maggirwar, S.B., and Bambara, R.A. (2017). Deficiency in DNA damage response, a new characteristic of cells infected with latent HIV-1. Cell Cycle 16, 968–978.

Piret, J., and Boivin, G. (2016). Antiviral resistance in herpes simplex virus and varicella-zoster virus infections: diagnosis and management. Curr Opin Infect Dis 29, 654–662.

Pommier, Y. (2013). Drugging topoisomerases: lessons and challenges. ACS Chem Biol 8, 82–95.

Pourchet, A., Copin, R., Mulvey, M.C., Shopsin, B., Mohr, I., and Wilson, A.C. (2017). Shared ancestry of herpes simplex virus 1 strain Patton with recent clinical isolates from Asia and with strain KOS63. Virology 512, 124–131.

Simsek, D., Brunet, E., Wong, S.Y., Katyal, S., Gao, Y., McKinnon, P.J., Lou, J., Zhang, L., Li, J., Rebar, E.J., et al. (2011). DNA ligase III promotes alternative nonhomologous end-joining during chromosomal translocation formation. PLoS Genet 7, e1002080.

Sliter, D.A., Martinez, J., Hao, L., Chen, X., Sun, N., Fischer, T.D., Burman, J.L., Li, Y., Zhang, Z., Narendra, D.P., et al. (2018). Parkin and PINK1 mitigate STING-induced inflammation. Nature.

Smith, S., Reuven, N., Mohni, K.N., Schumacher, A.J., and Weller, S.K. (2014). Structure of the HSV-1 genome: manipulation of nicks and gaps can abrogate infectivity and alter the cellular DNA damage response. J Virol.

Suberbielle, E., Sanchez, P.E., Kravitz, A.V., Wang, X., Ho, K., Eilertson, K., Devidze, N., Kreitzer, A.C., and Mucke, L. (2013). Physiologic brain activity causes DNA double-strand breaks in neurons, with exacerbation by amyloid-β. Nat Neurosci 16, 613–621.

Szymonowicz, K., Oeck, S., Malewicz, N.M., and Jendrossek, V. (2018). New Insights into Protein Kinase B/Akt Signaling: Role of Localized Akt Activation and Compartment-Specific Target Proteins for the Cellular Radiation Response. Cancers (Basel) 10.

Terenzio, M., Schiavo, G., and Fainzilber, M. (2017). Compartmentalized Signaling in Neurons: From Cell Biology to Neuroscience. Neuron 96, 667–679.

Thellman, N.M., and Triezenberg, S.J. (2017). Herpes Simplex Virus Establishment,Maintenance, and Reactivation: In Vitro Modeling of Latency. Pathogens 6.

Vink, E.I., Lee, S., Smiley, J.R., and Mohr, I. (2018). Remodeling mTORC1 Responsiveness to Amino Acids by the Herpes Simplex Virus UL46 and Us3 Gene Products Supports Replication during Nutrient Insufficiency. J Virol 92.

Wan, G., Zhang, X., Langley, R.R., Liu, Y., Hu, X., Han, C., Peng, G., Ellis, L.M., Jones, S.N., and Lu, X. (2013). DNA-damage-induced nuclear export of precursor microRNAs is regulated by the ATM-AKT pathway. Cell Rep 3, 2100–2112.

Warren, S.L., Carpenter, C.M., and Boak, R.A. (1940). SYMPTOMATIC HERPES, A SEQUELA OF ARTIFICIALLY INDUCED FEVER: INCIDENCE AND C ASPECTS; RECOVERY OF A VIRUS FROM HERPETIC VESICLES, AND COMPARISON WITH A K STRAIN OF HERPES VIRUS. J Exp Med 71, 155–168.

Weitzman, M.D., and Fradet-Turcotte, A. (2018). Virus DNA Replication and the Host DNA Damage Response. Annu Rev Virol 5, 141–164.

Wheeler, C.E. (1975). Pathogenesis of recurrent herpes simplex infections. J Invest Dermatol 65, 341–346.

Wilcox, C.L., and Johnson, E.M. (1987). Nerve growth factor deprivation results in the reactivation of latent herpes simplex virus in vitro. J Virol 61, 2311–2315.

Wilcox, C.L., and Johnson, E.M. (1988). Characterization of nerve growth factor-dependent herpes simplex virus latency in neurons in vitro. J Virol 62, 393–399.

Wilcox, C.L., Smith, R.L., Freed, C.R., and Johnson, E.M. (1990). Nerve growth factor-dependence of herpes simplex virus latency in peripheral sympathetic and sensory neurons in vitro. J Neurosci 10, 1268–1275.

Wilkinson, D.E., and Weller, S.K. (2004). Recruitment of cellular recombination and repair proteins to sites of herpes simplex virus type 1 DNA replication is dependent on the composition of viral proteins within prereplicative sites and correlates with the induction of the DNA damage response. J Virol 78, 4783–4796.

Wilson, A.C., and Mohr, I. (2012). A cultured affair: HSV latency and reactivation in neurons. Trends Microbiol 20, 604–611.

Wu, Z.H., Shi, Y., Tibbetts, R.S., and Miyamoto, S. (2006). Molecular linkage between the kinase ATM and NF-kappaB signaling in response to genotoxic stimuli. Science 311, 1141–1146.

Zeng, Z., Cortés-Ledesma, F., El Khamisy, S.F., and Caldecott, K.W. (2011). TDP2/TTRAP is the major 5’-tyrosyl DNA phosphodiesterase activity in vertebrate cells and is critical for cellular resistance to topoisomerase II-induced DNA damage. J Biol Chem 286, 403–409.

Álvarez-Quilón, A., Serrano-Benítez, A., Lieberman, J.A., Quintero, C., Sánchez-Gutiérrez, D., Escudero, L.M., and Cortés-Ledesma, F. (2014). ATM specifically mediates repair of doublestrand breaks with blocked DNA ends. Nat Commun 5, 3347.

